# V1 neurons encode the perceptual compensation of “false torsion” arising from Listing’s law

**DOI:** 10.1101/428441

**Authors:** Mohammad Farhan Khazali, Peter Thier

**Author notes:** Address correspondence to Peter Thier, Hertie Institute for Clinical Brain Research, Dept. Cognitive Neurology, Hoppe-Seyler-Str. 3, 72076 Tuebingen, Germany.

## Abstract

We try to deploy the retinal fovea to optimally scrutinize an object of interest by directing our eyes to it. Horizontal and vertical components of these fixation eye movements are determined by the object’s location. However, fixation eye movements also involve a torsional component, which according to Listing’s law is fully determined by the 2D eye position acquired. According to Von Helmholtz knowledge of the torsion provided by this law alleviates the perceptual interpretation of the image tilt that changes with fixation, a view supported by psychophysical experiments he pioneered. We address the question where and how Listing’s law is implemented in the visual system and we show that neurons in monkey area V1 use knowledge of torsion to compensate the image tilt associated with specific eye positions as set by Listing’s law.

## Introduction

We explore a visual scene by a sequence of saccades and brief moments of fixation. According to Listing’s law^1^, the movements of our eyes from one position to the next are rotations around an axis that lies in a plane, whose orientation relative to the head is determined by the starting position of the eyes^2^. In the specific case of the eyes starting from straight ahead, i.e. from the primary position, this plane has a frontoparallel orientation and is usually referred to as Listing’s plane. The consequence of this principle is the emergence of an amount of torsion of the eye around the line of sight (*false torsion*) that is a function of the horizontal and vertical deviation from straight ahead, independent of the starting position of the eyes^1–3^. The amount of *false torsion* is zero for fixation positions on the horizontal and vertical head-centered meridians and significant for positions on or close to the diagonals where it grows with eccentricity (**Supplementary Fig. S1**). For instance, fixating at (35°, 35°) will be accompanied by around 15° of clockwise *false torsion* relative to the primary position. Although the *false torsion* will inevitably rotate the image on the retina and thereby rotate the world vertical relative to the retina vertical, we do not perceive a tilting world and in general, are not aware of any retinal consequences of this eye torsion^2,4^. Based on afterimage experiments^2^, Hermann von Helmholtz, to whom we owe the detailed elaboration of Listing’s law, could show that this is a consequence of a perceptual reinterpretation of the image orientation, based on knowledge of the amount of *false torsion*^4–7^. It is the integration of this torsion prior into the perceptual interpretation of retinal images that allows us to warrant the stable perception of object and world orientation when exploring visual scenes. Here we show that the perceptual interpretation is taking place at area V1 as its neurons preferred orientations and receptive field (RF) shifts with respect to the retina in order to compensate for *false torsion* in a way that explains the monkey’s recorded perception.

## Results

### Monkeys’ perception accounts for false torsion

We wanted to know where and how information on *false torsion* is integrated into the processing of the retinal image in the visual system at the level of single neurons. To answer this question, we resorted to macaque monkeys as they explore scenes like humans shifting fixation from one position to the next, therefore burdening also their visual system with *false torsion-based* image rotations^8–11^. In order to clarify if they are able to perceptually compensate the resulting retinal image tilt like humans, we trained one of the two monkeys (M1) on a two alternative forced choice task in which he had to decide if a slim line (14° visual angle length centered on the fixation dot) was tilted clockwise (*cw*) or counter-clockwise (*ccw*) relative to the external world-centered vertical with the head immobilized upright via an implanted head post (**Fig. 1A**). The line whose tilt was randomly selected from a set of tilt angles (method of constant stimuli) was present during 1 sec-periods of stable fixation (eye position within a window of 1° × 1°). After this period both the line and the fixation dot disappeared and two response targets, one on the left, the other one on the right side appeared, prompting the monkey to saccade to the right in the case of perceived *cw* tilt and to the left for perceived *ccw* tilt. In order to facilitate understanding the behavioral requirements, we added a visual reference line aligned with the world vertical in the early phase of the behavioral training (for details see Online Methods). In these “training trials”, the monkey quickly attained a high and reliable level of performance: whenever the tilt exceeded 6°, the monkey reported the correct tilt direction in >90% of the trials and responded at chance level (50%) when the line tilt was around zero (**Supplementary Fig. S1B**). This chance level point—the point of equal selection (*PES*)—served as a measure of the monkey’s subjective visual vertical (*SVV*) in the actual experiment starting after several weeks of intensive training. Here we stopped presenting the visual reference line, assuming that the monkey would now use an internal reference corresponding to his subjective vertical. The results described below support the conclusion that this was indeed the case. The monkey was given a reward whenever he correctly reported the line tilt for angles exceeding 6° in either direction but was rewarded at random, independent of his perceptual decision, for angles smaller than 6°. The latter trials (“test trials”) were always few (10–5%) and randomly interwoven into the other trials. *SVV* measurements were based on the test trials and were carried out for three different gaze positions: central (0°, 0°), upper right (20°, 20°) and upper left (−20°, 20°) with respect to straight ahead. These gaze positions were acquired by presenting the fixation point on one of three monitors, positioned tangentially on a virtual spherical surface (radius 107 cm) centered on the midpoint between the monkey’s eyes (**Fig. 1A**). When M1 gazed straight ahead, the *SVV* was 89.37° (95%, confidence bounds (conf. b.) 89.36, 89.39), i.e. deviating only slightly, yet significantly (p<0.005, t-test) from the world vertical in a *cw* direction (**Fig. 1B**). When the gaze was shifted to the upper left, the eyes rotated significantly by 5.1° (95% conf. b. 5.08, 5.12, pooled across days and both eyes, **Supplementary Fig. S3**) ccw, data pooled across days and eyes; see Online Methods for details on the eye torsion measurements) about the gaze axis relative to the eyes looking straight ahead (p<0.05, t-test). The *SVV*, on the other hand, displayed a comparatively minor, yet significant change (p<0.05) by 1.59° *ccw* to 90.96° (95% conf. b. 90.93, 90.97). If the *SVV* had been referenced to the retina, one would have expected a rotation of the *SVV* by an amount corresponding to the eye rotation and no change whatsoever in the case of a head- or world-centered reference system. This means that the *SVV* stayed closer to the world-centered vertical, considering that 65% of the eye rotation had been taken into account to compensate for the retinal image torsion due to the eye torsion. When the gaze was shifted to the opposite side, i.e. to the upper right, the eye torsion amounted to 5.5°(95% conf. b. 5.47, 5.53, pooled across days and both eyes) *cw* relative to straight ahead with respect to the central fixation (p<0.005, t-test). Again the *SVV* changed very little, yet significantly by 0.15° *cw* to 89.22° (conf. b. 89.2, 89.25 (p<0.005 t-test) relative to the gaze straight baseline). This again demonstrates a substantial, albeit not complete (97%) compensation of the perceptual consequences of the eye torsion for the *SVV*. Finally, when directly comparing the two eccentric gaze positions, the average difference in *SVV* orientations amounted to 1.74° (conf. b. 1.56, 1.92), whereas the average difference in eye torsion was 10.2° (conf. b. 10.16, 10.24). In other words, the perceptual system compensated on average 82% of the tilt of the *SVV* due to the differences in eye torsion associated with the changes in fixation positions. This amount of compensation is very similar to the one reported for humans^4^ and in humans large enough to avoid awareness of eye torsion-induced image tilt under natural viewing conditions.

**Figure 1.**
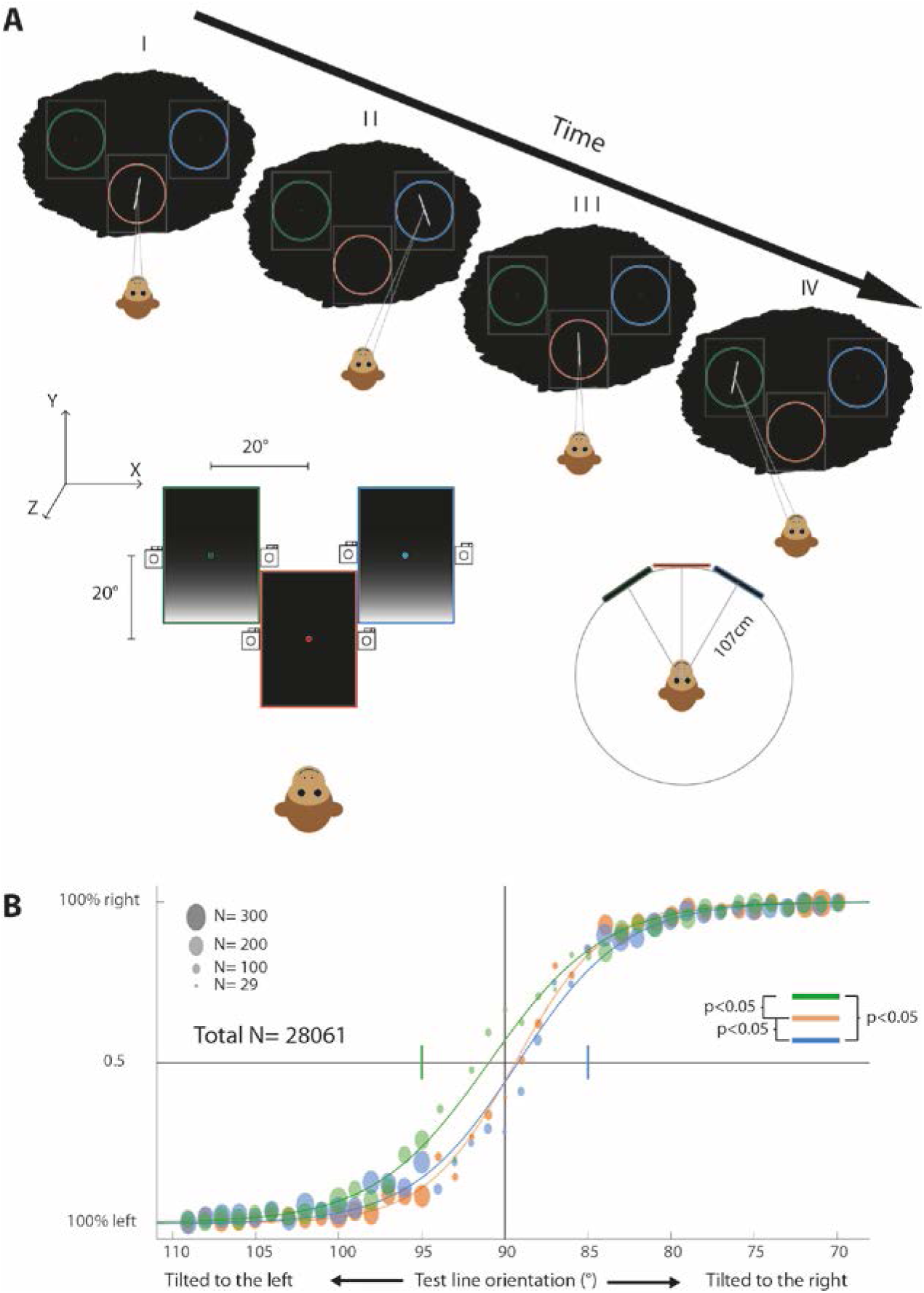
Assessing the subjective visual vertical of monkeys for different gaze directions. **A** Lower illustrations: The monkey fixated a target presented in the center of three monitors arranged tangentially on a 107 cm radius sphere with the monkey in the center of the sphere. One of the monitors was positioned straight ahead (0°, 0°) and the two others in the upper right (20°, 20°) and left (−20°, 20°) position respectively. Each monitor was equipped with two cameras, one facing the left eye, the other one the right eye, providing eye snap shots used to determine the amount of *false torsion*. Upper illustration: sequence of psychophysical tests in a typical daily experimental session probing the monkey’s *SVV*. The fixation target appeared on one of the three monitors and the monkey had to report the orientation of the test line relative to a vertical reference line (not shown) or relative to his internal reference (i.e. the *SVV*) by making an indicative saccade at the end of a trial. At the end of a block of trials the target and the test line jumped to another monitor. Note that a black circular aperture (indicated by the colored circles) covered the monitor edges (indicated by the grey rectangle) in order to eliminate any external orientation cues**. B** Plot of monkey M1’s decisions on the orientation of the test line (in the absence of a visual reference) as function of the true test line orientation. The plot is based on all daily experimental sessions including around 28,000 trials. The colors distinguish data for the three monitors/gaze directions (green: upper left monitor, blue: upper right, red: central). The chance level point (50% right vs. left choices, the point of equal selection (*PES*)) was taken as a measure of the monkey’s subjective visual vertical (*SVV*). Note that the *SVV* for the upper left monitor is shifted slightly in a counter clockwise direction, and for the upper right monitor in a clockwise direction with respect to the *SVV* for the central monitor. The *SVV* differences between monitors/gaze directions are minimal compared to the differences in *false torsion* whose range is pegged by the short vertical green and blue lines indicating eye torsion for the upper left and the upper right monitor respectively.

### V1 neurons compensate diversely for false torsion

After having established that monkeys’ perception of image tilts due to *false torsion* corresponds to that of humans, we embarked on a search for its neuronal correlate. In order to explain the percept, the neuronal correlate should reflect the extensive, yet not complete transformation of visual orientation responses from retinal into world-/head-centered coordinates based on the integration of information on *false torsion*. We assumed that this integration could take place already at the level of area V1 in view of the exquisite sensitivity of V1 neurons to the tiny changes in object image orientation and position as resulting from *false torsion*.

To critically test this idea, we compared the receptive field (*RF*) positions and preferred orientations of V1 neurons for at least two and in many cases for all of three gaze orientations studied in the psychophysical experiment on the *SVV* of M1. This comparison was carried out in M1 and a second monkey M2, for whom we did not attempt to collect perceptual data.

**Figures 2A** and **2B** depict the orientation tuning curves of two exemplary V1 neurons that represent the diversity of responses to *false torsion*. Tuning curves were obtained by flashing Gabor gratings centered on the neuron’s *RF*, whose orientation was varied at random in steps of 2°. The size, spatial frequency and duty cycle of the grating stimuli were adapted to the needs of the individual neuron (see Online Methods for more details). The tuning curves in **Fig. 2A, B** are plotted in a head-centered frame of reference (*FOR*) on the left and in a retina-centered *FOR* on the right. The neuron shown in **Fig. 2A** exhibited a tuning curve that stayed put when plotted in retinal coordinates, independent of the difference in eye torsion. In other words, this neuron showed behavior in accordance with the standard notion of retina-centered coding in V1. However, the neuron depicted in **Fig. 2B** presented a tuning curve that shifted on the retina by 9.42° (conf. b. 10.75, 8.09), i.e. relatively close to the *false torsion* difference measured in this particular session, which amounted to 11.7° (conf. b. 11.55, 11.85). Correspondingly, when plotted in head-centered coordinates, the orientation tuning curves did not change much with *false torsion*. We analyzed the dependence of the orientation preferences on the amount of *false torsion* associated with eccentric gaze in 129 neurons, 63 from M1 and 66 from M2, all having *RFs* in the lower left visual quadrant and most of them located in layers 2 and 3, based on the criteria of Snodderly and Gur^12^ (see **Supplementary Fig. S4B, C**). In order to quantify changes in orientation preferences prompted by eye torsion we calculated an orientation updating index (*OUI= - orientation change/eye torsion change*) defined by the inverted difference between preferred orientations in head-centered coordinates for two gaze directions divided by the associated amount of eye torsion change between the two. The angular change in preferred orientation from one gaze orientation to another one was obtained by determining the location of the peak of the cross correlation function between the individual tuning curves plotted in head-centered coordinates. The *OUI* is close to 0 for a neuron, which—like the one shown in **Fig. 2B**—is able to stabilize the orientation tuning curve relative to the head by fully compensating changes in eye torsion. It will be -1 for a neuron like the one shown in **Fig. 2A**, lacking any compensation of eye torsion, encoding the visual world in retina-centered coordinates. *OUIs* larger than zero reflect overcompensation of eye torsion and *OUIs* smaller than -1 erroneous shifts of the orientation preference in a direction opposite to the one allowing compensation of eye torsion. **Figure 2C** plots the distribution of the *OUI* for the 129 neurons recorded from M1 and M2. Pooling the data obtained from the two monkeys seemed justified as individual distributions did not differ significantly (p=0.43, Wilcoxon rank sum test). The pooled distribution is quite broad with two peaks, a larger one close to zero and a second one close to -1, well fit by a combination of two Gaussians peaking at 0.004 (conf. b -0.02, 0.03) and at -0.8102 (conf. b. -0.9069, -0.7135) respectively (see Fig. 2 for details on the statistics). This bimodality could indicate the presence of two distinct neuronal populations in V1, a larger one preferring a head-centered *FOR* and a smaller one preferring retina-centered coding. If the former were derived from the latter by integrating information on eye torsion, one might assume that it should be the former one that is responsible for the percept. Given the fact that the former mode peaks very close to zero one would expect a perfect percept, i.e. a *SVV* not deviating from the head-centered vertical. However, this is not the case as the *SVV* measured in M1 reflects a perceptual undercompensation of eye torsion. Actually, a comparable perceptual undercompensation would be expected under the assumption of joint action of all neurons contributing to both modes. The population mean of the individual *OUIs* amounts to -0.32 (conf. b.-0.47,-0.17) in monkey M2, indicating substantial undercompensation and to -0.23 (conf. b.-0.31,-0.15) in monkey M1 for whom psychophysical estimates of the *SVV* are available. We calculated a measure of perceptual compensation for this latter monkey, the subjective visual vertical updating index *SVV_UI*, given by the deviation of the *SVV*, for eccentric gaze, from the *SVV* at straight-ahead gaze divided by the amount of *false torsion*. The latter measure had a value of -0.172 (conf. b. -0.193, -0.151), not significantly different from the *OUI* of M1 (p=0.8) and the *OUI* of both M1 and M2 pooled (p=0.35). In other words, the features of the perceptual updating of the *SVV* may be understood as reflections of changes in the orientation preferences of an undifferentiated sample of 129 V1 neurons predominantly collected from layers 2 and 3. We arrived at the same conclusion when comparing normalized population orientation curves for the three gaze directions. Normalization was achieved by first scaling discharge rates for the various grating directions relative to the discharge rate for the preferred direction set to 1. Next, we rotated the orientation tuning function of a given neuron obtained for the central monitor such as to align the preferred direction of this neuron with the head-centered horizontal and rotated the same neuron’s tuning functions for the eccentric monitors by the same amount. **Figure 2D** depicts the resulting normalized population orientation tuning functions for the three monitor positions on top of each other in a head-centered *FOR*. All three tuning curves are well aligned with the head-centered horizontal, a fact that supports the notion of a population-based encoding of visual orientation in a head-centered *FOR* in V1, independent of gaze direction. The same conclusion is suggested by a mean *OUI* of -0.28 (conf. b. -0.31, -0.25), based on the average of the three pairwise comparisons of the *OUIs* of the normalized population tuning curves for the three gaze directions (see **Fig. 2D**, upper right).

**Figure 2.**
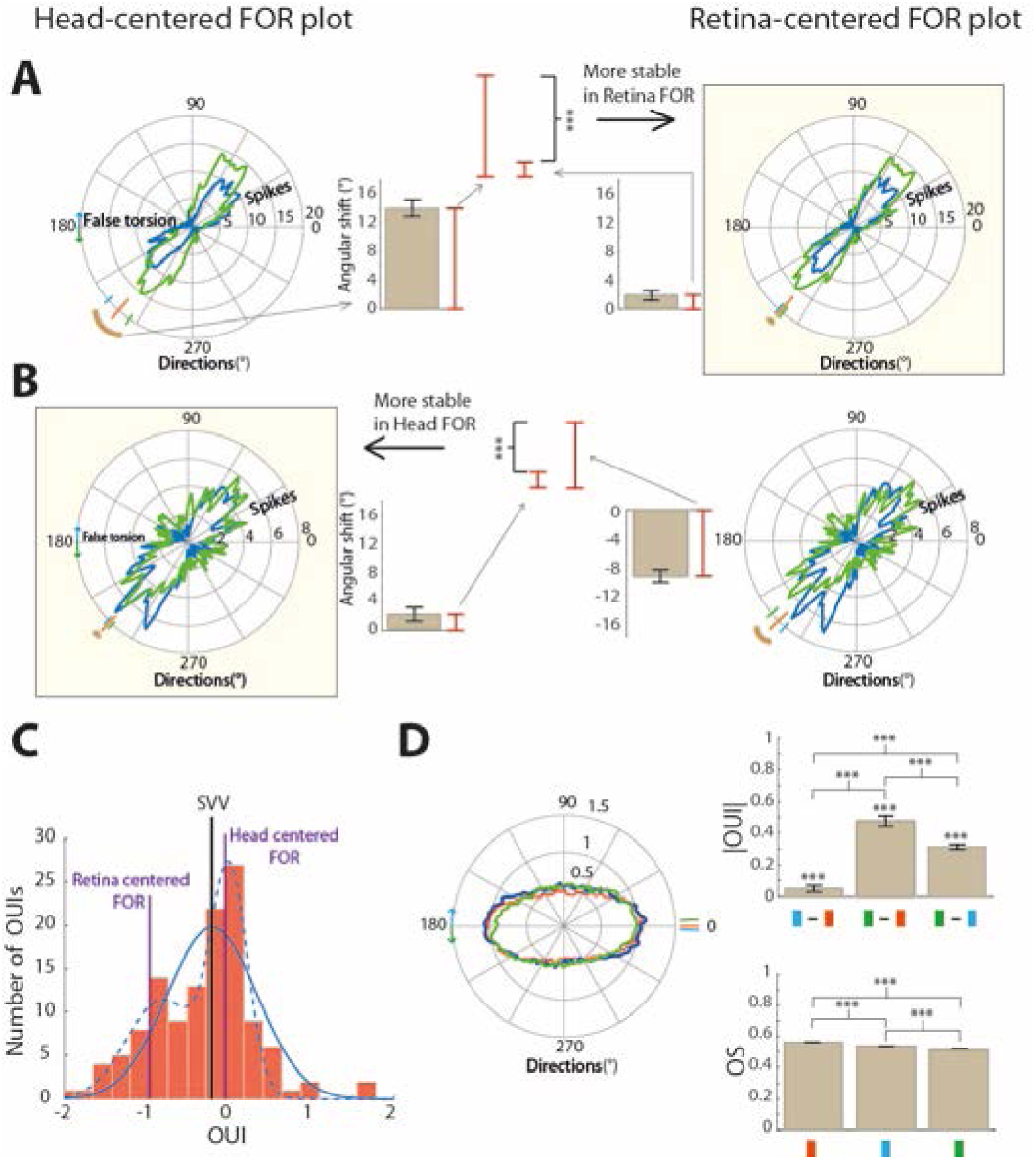
Diverse effects of *false torsion* on orientation tuning curves of V1 neurons. **A** and **B** show superimposed orientation tuning curves of two exemplary neurons obtained with the monkey fixating either on the upper left monitor (data in green) or the upper right one (data in blue). The angular axis represents the grating orientation and the radial axis the discharge rate evoked. On the left side we plot the orientation tuning curves relative to the monitor, i.e. in a head-centered frame of reference (*FOR*) and on the right in a retina-centered *FOR*. The green and blue arrows in the head-centered *FOR* plots mark off the difference in eye torsion between fixation on the upper left and right monitors. The small green, red and blue lines perpendicular on the outer circle of the polar plots indicate the preferred grating orientation for the upper left, middle and upper right monitor in the respective frame of reference (head-centered *FOR* on the left, retina-centered *FOR* on the right). The preferred orientation is perpendicular to the preferred direction of grating movement. The brown arc segments represent the angular difference between preferred orientations for the two. This difference is also reflected in the bar plot (mean ± standard error of the mean (se)) next to the orientation tuning plot. The orientation preference of the neuron shown in (**A**) is more stable in a retina-centered *FOR* as the angular difference between preferred orientations is much smaller for this *FOR* than for the head-centered one. The reverse is true for the neuron shown in (**B**). **C** Histogram of the orientation updating indices (*OUI*) of all neurons from both monkeys. Fully head-centered neurons have an *OUI* of 0, those which are retina-centered one of 1. Note that each *OUI* represents pairwise comparisons between the tuning curves associated with two different monitors. Hence, if a neuron were tested for all three monitors (yielding three tuning curves) it would contribute to the histogram with three individual *OUI* values. The solid and dashed blue lines depict the fits with one or two Gaussian functions respectively. The two Gaussian fit (r^2^=0.97 rmse=0.08) was better than the one Gaussian fit (r^2^=0.86 with rmse=1.57). The solid black vertical line indicates the averaged *SVV_UI* change based on pooling the data for all three monitors collected for M1. **D** The left panel shows normalized population orientation functions based on pooled data from both monkeys. The three superimposed functions represent the normalized population responses obtained when probing the orientation sensitivity with fixation on the three monitors with the curve for the central monitor serving as reference (see Online Methods for details). The upper right panel is a bar chart comparing the mean absolute *OUIs* of the normalized population tuning curves for the three gaze directions. The lower right panel depicts the mean orientation selectivity (*OS*) of the neurons for fixation on the three monitors. Note that the orientation selectivity is significantly lower for eccentric fixations (*** = p<0.001, t-tests). The error bars indicate se.

### V1 activity reflects perception of orientation

On closer inspection, it is noticeable that the population curve for gaze straight ahead is slimmer than the curves for the two eccentric gaze directions. This impression is supported by a quantification based on an orientation sensitivity index *OS*, which compares the discharge strength for the preferred orientation relative to the orthogonal one (*OS=* (*preferred orientation-orthogonal orientation*)*/* (*preferred orientation + orthogonal orientation*). The *OS* was indeed significantly larger for straight ahead (OS=0.56 ± 0.003 mean ± se) than for the two other gaze directions (upper right OS=0.54 ± 0.0024, and upper left OS=0.52± 0.0021). Basically, the same result was obtained if only neurons from M1 were considered (**Fig. 3B**). Assuming that orientation judgments are based on a population vote of all neurons in monkey M1, less sharp orientation tuning of the population responses for eccentric gaze should translate into less certain orientation judgments. This is exactly what the perceptual data for monkey M1 exhibited: as shown in **Fig. 3A** the steepness (*a*) of the psychometric function quantifying M1’s perceptual decisions around the point of *SVV* was significantly less (p<0.001) for eccentric gaze than for gaze straight ahead. The notion of a close link between the precision of orientation judgments and the properties of neurons is also supported by the fact that the changes of *OUIs* between monitor pairs (e.g. comparison of mean *OUIs* between the upper left and central monitor) matched the changes of the corresponding *SVV_UIs* very well (**Fig. 3C**).

**Figure 3.**
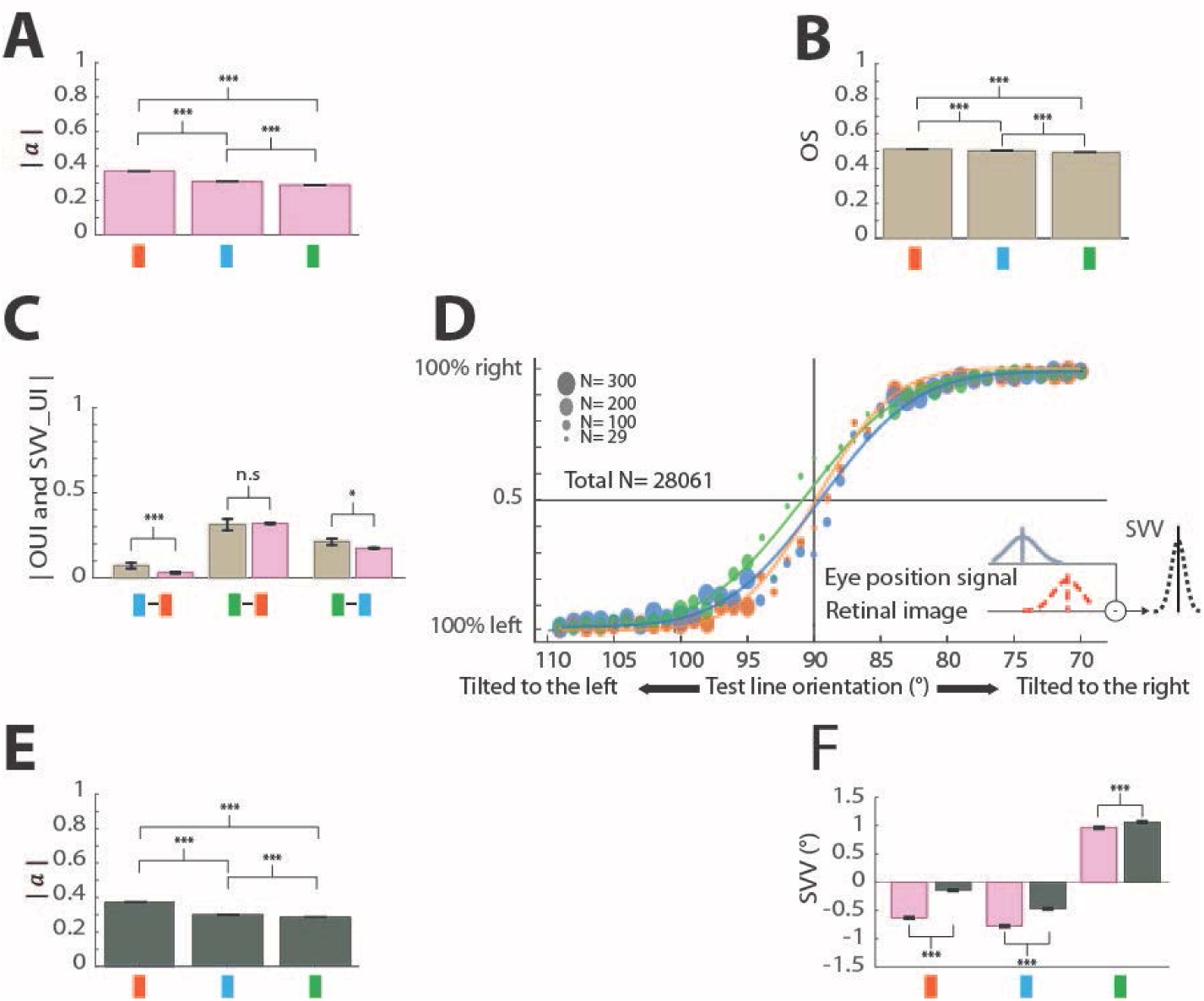
Explaining the *SVV* based on the integration of a retinal orientation signal and an eye torsion prior. **A** Bar chart comparing the absolute value of the *a* parameter of the psychometric functions (see Materials Methods) fitting the orientation decision data pooled over all experimental sessions, a measure of the precision of orientation judgments for the three gaze directions tested. Note that the *|a|* measure for gaze straight ahead was significantly larger than for the fixation on the two eccentric monitors (0.37±8.96·e-4, mean ±se for the central monitor; 0.31±8.6·e-4 for the upper right monitor and 0.29±46·e-4 for the upper left monitor)**. B** Bar chart of the mean orientation selectivity (*OS*=0.51±0.126·e-3 mean±se for the central monitor, 0.50±0.146·e-3 for the upper right monitor and 0.49±0.146·e-3 for the upper left monitor) of all neurons recorded from M1 and tested for fixation on the three monitors. **C** Bar chart comparing the mean *OUIs* (brown) of the normalized population tuning curves for the three gaze directions in M1 and the *SVV_UI* (pink). Note that we are plotting the absolute values of the *OUIs* and *SVV_UI*. D Fits of model function to the psychophysical data on the *SVV*, also presented in **Fig. 1B**. The green, red and blue colors distinguish data and fitted functions for fixation on the upper left, the central and the upper right monitor respectively. The inset tries to capture the model structure. **E** Bar chart summarizing the *|a|* values of the psychometric functions (means ±se) predicted by the model for the three fixation conditions (red for the central monitor, blue for the upper right and green for the upper left monitor). Note that the predicted *|a|* shows a dependence on the monitor position similar to the position dependence of the measured *|a|* (0.38±3.36·e-12, 0.30±1.26·e-50 and 0.29±16·e-50) shown in (**A**). **F** Bar chart indicating the measured *SVV* (pink) and the *SVV* estimated by the model (dark grey) for the three gaze directions. Note that the model was able to predict the direction and relative amount of error of *SVV* across monitors quite well (*** = p<0.001, t-tests).

We explored the gaze direction-dependent changes of the neuronal responses and their relationship to the perceptual judgments further by modeling perceptual judgments based on the assumption that they reflect the integration of information on visual orientation and on eye torsion. The model (see Online Methods for details) assumes that information on visual orientation is originally available in a retinal *FOR*. As the visual signal reflects the collective vote of a number of orientation-selective neurons tuned to individually different preferred orientations, the noise level should not depend on the specific orientation preference, for the sake of simplicity ignoring here the complication of the oblique effect^13–15^. On the other hand, eye torsion-associated noise should grow with torsion as torsional information is assumingly conveyed by neurons encoding eye torsion in a monotonic format^16,17^. This two component model, fitted to the three psychometric curves, was able to reproduce the observed *SVV* associated with the three fixation positions very well (r^2^ = 0.99; root mean of squared error (rmse) = 0.006; **Fig. 3D**, **E** and **F**), including the perceptual undercompensation of *false torsion* and the lower *a* of the psychometric functions for the eccentric fixation positions. Assuming that the perceptual judgments are based on a V1 population vote, we suggest, that the increase in eye position-dependent noise assumed in the model explains the poorer orientation sensitivity of the population response for eccentric gaze directions. The fit also accommodates the tilting of the *SVV* by about 0.15°cw relative to the true gravitational vertical due to a torsional bias of 0.53°cw during straight-ahead fixation (**Fig. 3F** and **Supplementary Fig. S9**), a perceptual bias that also contributes to the perceptual judgments for the eccentric gaze directions (gaze upper left: 1.06°ccw, gaze upper right: 0.47°cw). Subtracting the eye torsional bias leads to tilts of the *SVV* that become much more mirror symmetric relative to the true gravitational vertical.

### V1’s RFs account for false torsion

Torsional eye movements not only rotate the object image relative to the retinal meridian but also translocate the image if located outside the center of rotation. In order to warrant spatial stability, the *RFs* underlying the representation of the object should move relative to the retina in a fully compensatory manner. For 99 of them, 52 from M1 and 47 from M2, we were able to obtain detailed *RF* maps resorting to a reverse correlation approach (see Online Methods) not only for the straight-ahead eye position but also for at least one of the two eccentric gaze directions, in many cases for both (**Supplementary Fig. S4A**; **Supplementary Fig. S10A** and **B**). We quantified how much the angular position of *RFs* changed with eye torsion by calculating a *RF* updating index (*RFUI = - RF angular position change/eye torsion change*). A *RFUI* of -1 indicates retina-centered *RFs* and a *RFUI* of 0 head-centered coding, i.e. an updating of receptive field position by fully considering the amount of gaze direction-dependent eye torsion. The resulting distribution of *RFUI* (Fig. 4) is broad with most values located between *RFUI* of 0 and -1. However, rather than being uniform, it exhibits two distinct modes, well fitted by a sum of two Gaussian models (see the figure legend for details), a smaller mode peaking close to 0 and a larger one, peaking at -0.66. Whereas the former can be associated with head-centered coding, the latter mode is intermediary between the two frames of reference, formed by neurons that exhibit varying degrees of partial consideration of eye torsion. Note that these results cannot be influenced by possible differences in the quality of fixation on the three monitors as there were none: when plotted in monitor coordinates the mean fixation positions for the three monitors were the same for eye position data sampled together with the neuronal data (Kruskal Wallis ANOVA with the factor gaze direction, p= 0.12 and p=0.39 for horizontal eye position for M1 and M2 respectively, and p=0.11 and p=0.39 for vertical eye position for M1 and M2 respectively). Also the eye position variance for the three monitors did not differ significantly (p= 0.14 for M1 and p=0.37 for M2). Gaze direction-dependent changes in eye torsion affected the angular position of *RFs*, yet did not change their eccentricity. Accordingly, we did not find any change in mean eccentricity of *RF* locations with gaze direction (**Supplementary Fig. S11A**). Finally, gaze direction-dependent changes in eye torsion can be expected to affect the orientation preference and the angular position of its *RF* in a yoked manner. That this is indeed the case is supported by the fact that the amount of *RF* updating correlated significantly with the orientation preference shifts recorded from the same neuron (**Supplementary Fig. S11B**).

**Figure 4.**
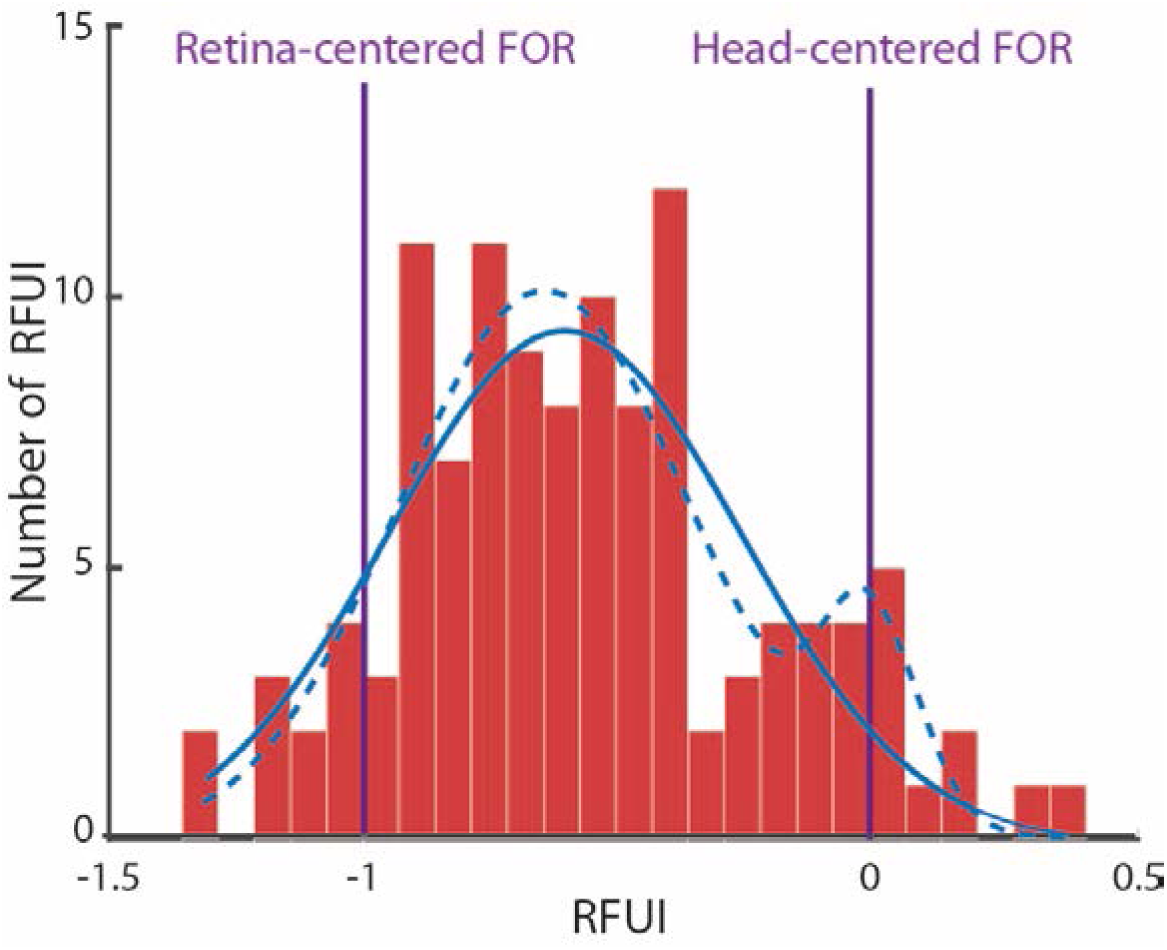
Impact of false torsion on the *RFs* of V1 neurons. Histogram of the receptive field updating index *RFUI= RF shift / torsional eye* shift of all neurons collected from M1 and M2. The two Gaussian fit explains the histogram better (r^2^=0.84 and rmse=1.8) than the one Gaussian fit (r^2^=0.69 and rmse=2.2). Note that individual neurons may contribute one or three *RFUIs*, depending on whether RF maps were obtained for two or three gaze directions.

## Discussion

Listing’s law alleviates the control of exploratory eye movements by reducing the degrees of freedom from 3 to 2 by making eye torsion dependent on the x-, y-eye position^1,2^. By the same token, it also alleviates the perceptual interpretation of retinal image orientation by constraining eye torsion—and consecutively the amount of image torsion relative to the retinal meridian—to just one value, firmly and reliably associated with a given x-, y-eye position. The prior knowledge of this value, the amount of *false torsion*, allows the transformation of the retinal image, tilted by a small, yet significant amount for eccentric gaze, into a torsion-invariant frame of reference. Our study shows that information on *false torsion* is taken into account at the level of area V1, allowing V1 to make possible a representation of object orientation that is independent of eye gaze direction-dependent torsion. Perceptually, *false torsion* is compensated largely, but not completely. The fact that the amount of perceptual undercompensation is predicted by the collective orientation vote of V1 neurons supports the notion that it is V1 that allows us to cope with the perceptual consequences of Listing’s law. The small amount of undercompensation of *false torsion* remains unnoticed outside the laboratory, i.e. in the absence of careful psychophysical measurements, and is therefore tolerable. A simple Bayesian model of orientation judgments, integrating information on visual orientation and *false torsion* suggests that the advantage of tolerating this small amount of undercompensation might be to limit the impact of eye position-dependent noise, more and more influencing perceptual judgments for larger amounts of eye torsion. *False torsion* not only influences image orientation but also image position on the retina. As we are lacking reliable information on the influence of *false torsion* on the perception of image position, it must remain open, if the *RF* position shifts observed are able to explain the percept. As the population data indicate that torsion-induced changes in object position are on average compensated only about half, one might also expect that the degree of perceptual position invariance may be substantially less than the perceptual orientation invariance. On the other hand, the evidence for a subgroup of neurons with almost perfect eye torsion invariance of *RF* positions may on the contrary suggest perfect torsion invariance, assuming that only this subgroup underlay the percept. However, this does not seem to be likely when considering that also orientation judgments were much better explained when considering the whole population of neurons, not only those found in the distribution mode, comprising largely torsion-invariant neurons. Actually, the scarce perceptual data on humans might be taken to suggest that subjects partially misjudge the position of a flashed probe with respect to the head plane during eccentric fixation^6^, supporting the assumption of a collective neuronal vote.

The fact that V1 takes information on eye torsion into account in order to compensate the consequences of Listing’s law for the perception of object orientation and position does not necessarily imply that V1 encodes visual information in a fully head-centered frame of reference, i.e. also compensating the horizontal and vertical deviations of the eyes relative to the head. *False torsion* due to Listing’s law is not the only form of eye torsion whose perceptual consequences are compensated by V1. Another form is ocular counter-roll evoked by tilting the head about the roll axis. A specific subset of V1 neurons uses information on counter-roll to render visual orientation and position in a head-centered *FOR*, thereby helping to establish a world-centered representation of the visual world, invariant to roll tilt of the head and body^18^. As ocular counter-roll is small, its contribution to a tilt-independent percept of the vertical is negligible. In fact, the generation of a tilt-invariant representation would be simplified considerably if the ocular counter-roll response to head tilt would be simply vetoed, making the consideration of eye torsion dispensable. Why is this not the case? The answer may be that perfect vetoing of counter-roll, a phylogenetic vestige of lateral eyed ancestry, may simply be unnecessary. Counter-roll is reduced by just an amount needed to make use of the machinery that is in any case needed to cope with the consequences of *false torsion*.

## Online Methods

### Subjects

Two adult male rhesus monkeys (Macaca mulatta) took part in this study (M1, 10 years old, M2, eight years old). The experiments were approved by the local authorities in charge (Regierungspräsidium Tübingen and Landratsamt Tübingen), conducted in accordance with German and European law and the Guidelines of the National Institutes of Health for the Care and Use of Laboratory Animals and carefully monitored by the veterinary service of Tübingen University. A magnetic scleral search coil was implanted into the right eye to record 2D eye position and a titanium head post to painlessly immobilize the head during experiments. We refrained from trying to document also torsion with search coils as we were concerned that the much bulkier 3D search coils might have a mechanical impact on the torsional eye position associated with eccentric fixation. Therefore, torsion was measured by comparing video images of the eyes acquired during stable fixation of the target presented in the center of the monitors. Six cameras were used, two for each monitor, in each case one for each eye, to guarantee an optimal alignment with the line of gaze of both eyes when the monkey fixated the center of a particular monitor, thereby minimizing image distortion. We implanted a circular titanium chamber over occipital cortex for electrophysiological recordings. All surgical procedures were conducted adopting aseptic techniques under adequate anesthesia consisting of isofluorane supplemented with remifentanil (1–2 μg/kg/min). All relevant physiological parameters such as body temperature, heart rate, blood pressure, pO_2_, and pCO_2_ were continuously monitored. Postoperatively, buprenorphine was given until no sign of pain was left. Animals were allowed to fully recover before starting the experiments.

### Single-unit recording

Extracellular action potentials were recorded with commercial glass-coated microelectrodes (Alpha Omega Engineering, impedance at 1 kHz: 0.5–1 MOhm). We inserted the electrode through the intact dura. We stopped advancing the electrode when it reached the brain and left it at that position for around 30 minutes to give the tissue a chance to relax after the penetration. Thereafter, we advanced very slowly (1 mm/h) to give the tissue sufficient time to adjust to the pressure exerted by the electrode. The position of a neuron relative to the top of cortex was estimated as the difference between the electrode position when recording the neuron’s action potential and the electrode position at which we observed the last neural activity when later extracting the electrode again. This estimate, the level of spontaneous activity of neurons at a given level, spike morphology, the preponderance of orientation versus direction selectivity and other criteria suggested by Snodderly and Gur^12^ were used to estimate the layer of a recorded neuron. Spikes of well-isolated single neurons were discriminated online by real-time sorting software based on template matching (Alpha Omega Engineering).

### Experimental setup and behavioral task

Three monitors (20° × 40°) were positioned tangentially on a virtual spherical surface (radius 107 cm) centered on the midpoint between the monkeys’ eyes. The central monitor was centered on the normal vector on the midpoint of the line, connecting the monkeys’ two eyes, taken as straight ahead (0°, 0°). The upper left monitor was centered at 20° left, 20° upward relative to straight ahead and the upper right monitor at 20° right and 20° upward. The alignment of the screens perpendicular to the line of sight was achieved by positioning a laser pointer in the monitor’s center with the laser oriented perpendicular to the screen such that the laser hit the midpoint between the monkeys’ eyes. This allowed us to ensure that the monitor surfaces were aligned tangentially on a virtual sphere centered on the midpoint between the monkeys’ eyes. Consequently, the surface of the central monitor was oriented parallel to the gravity vector, whereas the surfaces of the eccentric monitors, displaced up relative to the central one, were tilted by about 20°. In order to ensure that the vertical axes of the monitors were parallel to the gravity vector, we relied on information provided by an accelerometer system attached to the monitors (Analog Devices Inc., dual axis accelerometer ADXL203EB), allowing us to sense misalignments at a resolution of ±0.1°. Importantly, the alignment of monitors was checked and —if necessary—adjusted before each experiment. We measured the monkeys’ horizontal and vertical eye position using a self-made search eye coil system supporting a sampling rate of 1,000 samples/s. The search coil signal was calibrated using the known position of a white fixation target dot (diameter 0.48°) that appeared at random on the monitor in one out of nine positions defining a 3° × 3° grid centered on the monitor. Monkeys were asked to maintain fixation at each target for approximately 1 second in order to get a liquid reward and then to proceed to the next cued location. Targets were visible for 2 seconds and had to be fixated successfully at least three times. The data acquired was subjected to a regression analysis that considered linear, quadratic and mixed term dependencies in order to predict eye position based on the search coil voltage. In order to characterize the features of a neuron, after successful calibration the monkeys had to fixate the target (diameter=0.03) presented in the center of the central monitor for at least 10 minutes in total to allow us to map its *RF* and to assess its orientation preference. Fixation was considered acceptable if gaze stayed within a 1°-fixation window for 3 seconds. In this case the monkey was rewarded with a drop of juice or water, depending on the monkey’s preferences. Responses to visual stimulation were only considered if the aforementioned fixation requirement was met. Following testing visual stimuli on the central monitor, monkeys were required to fixate the same target presented on the eccentric monitors, one after the other in pseudorandom order across days in order to re-run the analysis of neuronal response features.

### Quality of fixation

Fixation noise, i.e. the variability in measured fixation position, the resultant of true biological fixation instability and electronic noise, was very low (mean ± standard deviation (stdv), 0.0058±0.0021° for M1 and 0.0063 ± 0.0019° for M2, pooled across all three monitors for periods excluding saccades and blinks; no significant differences between monitors; p>0.05). Considering the eye position during the stimulus presentation, both monkeys fixated on all monitors very well (see **Supplementary Table S1** for more details) with no difference in the average eye position and its standard deviation for fixation on the three monitors.

### SVV task

The visual stimuli were presented within circular dark apertures (diameter=16.5°), centered on the monitors, to prevent M1 from using the orientation of the monitor edges when judging the visual vertical (**Fig. 1A**). M1was trained to report the orientation of a tilted line with respect to the head/gravity vector. A trial started with a period in which M1 had to fixate a central dot (radius size 0.03°; fixation window 1°) for 800 ms, next a line (14° visual angle length, 0.8° width) with varying orientation was added, centered on the fixation dot. Continued fixation was required. After 1 second, this “test” line and the fixation dot disappeared and two targets, left and right respectively of the monitor center appeared for 700 ms asking the monkey to make a saccade to one of them, depending on the perceived orientation of the line. After that a bright, homogenous background was introduced for 500 ms to erase the afterimage of the line. The left target was associated with counterclockwise (*ccw*) line tilts relative to the gravitational vertical and the right one with clockwise (cw) tilts. In the beginning of the experiment, a reference line (16.5° length and 0.8° width) indicating the gravitational vertical was available throughout the trial to help the monkey to understand the relationship between the test line tilt direction and the two response targets. In these training sessions, in which stimuli were usually only presented on the central monitor (**Supplementary Fig. S2**), the monkey’s performance was excellent (>90%) for tilt angles of 6° or larger. For smaller tilt angles the accuracy of judgments decreased proportionally with the decrease in tilt angle until it reached the level of chance performance of (50%) for line tilts very close to zero (**Supplementary Fig. S1B**). At this point of subjective equality (*PSE*) the monkey obviously guessed, as he was no longer able to detect a consistent tilt. After several weeks of training we started presenting the test line without the reference line in test trials randomly interleaved with the training trials (see **Supplementary Fig. S2** for information on proportions) since the monkey seemed to have understood the need to base tilt decisions on a more general concept of the vertical. Whereas in the training trials the monkey was rewarded for correct choices, in the test trials a reward was given for correct choices in trials with tilt angles (relative to the gravitational vertical) exceeding >6° to either side as the monkey did not have any problems in detecting the true tilt direction, even though the reference line was missing. For smaller tilt angles <6°, rewards were provided at random on average in 50% of trials, independent of choices. There were always only a few trials with smaller tilts (10–15% of the test trials) in order to ensure that the monkey’s concept of an association of decisions (and rewards) on perceived line tilt would not be jeopardized. After several days, training trials could be fully omitted as M1’s responses to test trials suggested reliable perceptual reports of *SVV* (**Supplementary Fig. S2A**). At this point, we also started to test the monkey with stimuli presented on eccentric monitors. Responses were collected in blocks, usually starting with the stimuli presented on the central monitor, followed by the eccentric ones, pseudo-randomizing the order of the upper left and the upper right monitor (**Fig. 1A**, **Supplementary Fig. S2C**, and **D**). The *PSE* was determined by fitting a psychometric function 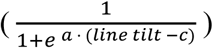 to the test trial data, pooled across all sessions, independent of whether interleaved training trials had been presented or not. In this function, *c* is the line tilt angle (*c*) for which the function predicted a *PSE* and the parameter *a* determines the steepness of the function. The *PSE* served as an estimate of the monkey’s subjective visual vertical (*SVV*) in the absence of a visual reference.

### Modeling the SVV

The model assumes that decisions on object orientation are based on the integration of information on eye gaze direction and the orientation of the object image on the retina and that both, visual and eye position-related signals are contaminated by noise. In the case of eye position, we suppose that noise increases with eccentricity considering the observation that eye position-related neurons typically exhibit a monotonic dependence of their discharge on eye position^16^. The signal on retinal image orientation is assumed to reflect the collective vote of a number of orientation selective neurons tuned to individually differing preferred orientations. The noise associated with information on retinal image orientation should be independent of the specific orientation as its extraction is based on dedicated neuronal filters, expected not to differ in terms of their signal-to-noise level. In other words, the vote on retinal orientation 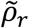 can be predicted by a Gaussian distribution with standard deviation *σ_r_* centered on the true retinal image orientation *ρ*. Note that— for the sake of simplicity—we ignore the well-established fact that neurons preferring orientations corresponding to the cardinal axes are overrepresented, therefore collectively offering a better signal-to-noise ratio, a fact that explains the psychophysical oblique effect^13–15^. The estimate of the retinal orientation is given by the likelihood function 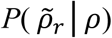.

The observer’s vote on the orientation of the object relative to the head 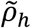 is the consequence of updating the retinal estimate of visual orientation 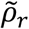 by the extra-retinal estimate of *false torsion 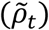*, given by the likelihood 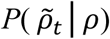. As said earlier, it assumes that eye torsion noise *σ_t_* increases monotonically with increasing deviation from zero eye torsion according to *σ_t_=σ*_0_ *+ σ*_1_ · *ρ* where *σ_0_* is the baseline noise associated with zero torsion and *σ_1_* the noise component scaled by eye torsion relative to zero torsion. The psychophysical data collected from M1 suggested that the torsional position associated with gaze directed at the straight-ahead monitor deviated slightly from zero torsion in the *ccw* direction. Hence, when looking at the left monitor the absolute torsional deviation from zero torsion was larger than when looking at the upper right monitor and, consequently, also the eye torsion-dependent noise component *σ*_1_·*ρ* larger for the left monitor. In other words, the torsion bias is taken into account by the model.

The estimate of the head-centered object orientation 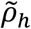 is given by the likelihood function 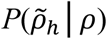 with 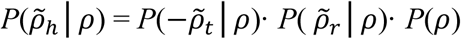

The negative sign inverting 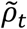 accounts for the fact that *cw* eye torsion causes *ccw* image rotation and vice versa. Note that *P*(*ρ*) =1 as we do not make any assumption on mean eye torsion. We take the maximum of the posterior probability *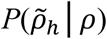* function as measure of object orientation relative to the head, i.e. (*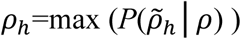*).

The relevant parameters were optimized by fitting the model to the three psychometric functions collected from M1, applying the Matlab function fminsearch, which is using the Nelder-Mead method, a multidimensional unconstrained nonlinear minimization method. The results are summarized in the legend of **Supplementary Figure S9**. We used a bootstrapping approach to estimate the reliability of the fit to the data. To this end, we fitted the model to randomly chosen 7,000 *SVV* measurements (with replacement) for each monitor. We repeated the procedure 100 times.

### Characterizing the RFs of V1 neurons

Testing individual neurons involved mapping their *RF* at high resolution, in most cases for all three gaze directions, but at least for two directions, while the monkey’s gaze stayed within a fixation window of 1° × 1°, centered on a 0.03° fixation target. *RFs* were determined by resorting to a reverse correlation approach^18^, which was based on probing parts of the visual field of interest with a flickering bright little squared dot (0.1 × 0.1°, contrast of 95%). The stimulus was on for 50 ms, followed by 50 ms of darkness and then reappeared in a new, randomly selected location, not overlapping with the preceding one, within a window of 3° × 3°, the latter centered on the expected location of the *RF*. Each element within the window was stimulated at least six times. The responses to the six runs of stimulus presentations were used to generate maps of the probability of dot presence within the window a certain time before the firing of an individual action potential (see **Supplementary Fig. S10A**). The temporal delay associated with the probability map that showed the highest peak when averaging the number of evoked action potentials across the y-axis was taken as the optimal latency for the stimulated neuron. The characterization of optimal latencies was carried out for each gaze direction without observing consistent differences in latencies between directions (1-way ANOVA, p>0.05). The probability map with the optimal temporal delay was taken as *RF* map, in which *RF* boundaries were delineated. After smoothing the *RF* map by adopting a 2D averaging filter with a 5-pixel radius, we applied Otsu’s method for global image thresholding, yielding binary maps. Next, we used the Matlab function bwboundaries.m to trace the exterior boundaries of the area containing “on” pixels. Note that the resulting contour lines representing *RF* boundaries were determined for visualization purposes only (as in **Supplementary Fig. S10A, B**) and not used to compare *RF* maps across conditions. To measure the amount of shift between *RF* maps for different gaze directions, the smoothed probability maps were cross-correlated using a normalized 2D cross-correlation approach and the shift measures obtained tested for significance by resorting to a bootstrapping method similar to Daddaoua and coworkers^18^. Briefly, since each part of the area of interest had been probed six times, we calculated six *RF* maps for each gaze direction. The resulting six *RF* maps were used to generate 1,000 *RF* maps by randomly (with replacement) selecting individual pixels from the individual six maps. This was carried out for each gaze direction. We then performed 1,000 2D cross-correlations between the resulting two *RF* map sets for each of the three pairs of gaze directions. We extracted the spatial coordinates associated with the maximum cross-correlation coefficient for each pair, allowing us to generate two distributions (each containing 1,000 values) for the x- and y-coordinates (**Supplementary Fig. S6D**). The means of these two distributions describe how much one should displace the *RF* map obtained for one gaze direction over the other gaze *RF* map to have both *RFs* overlapping. The means of the x- and y-distributions were taken as measures of the actual *RF* displacement. In order to test for significance, we generated noise distributions of the x- and y-components of spatial shifts by cross-correlating 1,000 pairs of *RF* maps for the same gaze direction and comparing the resulting distributions with the ones for the comparison of two gaze directions, using unpaired t-tests (p<0.05).

### Characterizing the orientation preference of V1 neurons and its dependence on gaze direction

Orientation tuning curves were assessed using drifting Gabor gratings present for 450 ms duration followed by 450 ms black screen. The grating appeared in an aperture of a diameter adjusted qualitatively for each neuron after being centered on the receptive field, the grating frequency and the speed of movement were selected to meet the spatial and temporal frequency requirements of the recorded neuron as suggested by prior qualitative testing with a limited set of test gratings. The orientation space was probed by moving the optimal grating in one of 180 directions differing by multiples of 2° drawn at random from trial to trial. Each movement direction was presented twice. To determine whether a given neuron was orientation or direction selective, we generated polar plots of discharge as function of grating movement direction by reverse correlating spikes with grating direction. We then applied a Fast Fourier Transformation (FFT) to the polar plot^19^. The zero order gain component of the resulting spectrum was taken as measure of the spontaneous activity, the first order component interpreted as the strength of the direction selectivity and the second order component as the strength of the orientation selectivity. A neuron was counted as orientation selective but not directional if the gain of the orientation component was greater than the gain of the direction component and, conversely, it was considered direction selective if the gain of the direction component was greater than the gain of the orientation component. Since circular means of the discharge rate cannot assess the preferred grating direction for orientation selective neurons as they have similar responses to two opposite directions of grating motion, we used a FFT to assess the preferred direction for all neurons. The preferred directions of both orientation and direction selective neurons were determined by fitting the three components of the spectral response function with the discharge rates. The peak of the fit to estimate the grating’s direction giving rise to the largest discharge rate was taken as the preferred direction, and the direction perpendicular to it was taken as the grating’s preferred orientation. After that, we tested the significance of the given neuron’s tuning function using a Rayleigh test as follows: If a neuron was classified as direction selective we applied a Rayleigh test directly to its tuning function. In case the neuron was classified as orientation selective we considered the two lobes of the polar plot separately and applied Rayleigh tests to both. All orientation selective neurons had both lobes passing the Rayleigh test. More than 90% of our recorded neurons were orientation selective based on the aforementioned criteria. We assessed the tuning curves of each neuron for at least two gaze directions if not three. In order to quantify the amount of shift of the preferred orientation from one gaze direction to a second one, we resorted to cross-correlation with bootstrapping statistics. The procedure started with increasing the number of repetitions for each condition. To this end, we reconstructed the tuning curves by combining data from two neighboring directions each, thereby increasing the number of trials available for each bin, now spanning 4°, to four. The responses in the four trials were used to generate four individual tuning curves. Based on these four original tuning curves we constructed 100 new ones by randomly (with replacement) combining bins from the four original tuning curves. These 100 tuning curves were generated for each gaze direction and served as the basis for the calculation of cross-correlation functions between orientation curves for the central and each eccentric monitor (giving 2 × 100 cross-correlation functions) and between the two eccentric monitors (yielding 1 × 100 cross-correlation functions). Similar to the statistical analysis of *RF* shifts, we extracted the angular coordinates associated with the maximum cross correlation coefficient, which resulted in 100 values for each comparison (**Supplementary Fig. S6C**). Note, that we deployed a standard 1D cross-correlation rather than a circular correlation in a polar coordinate system, as they both gave the same outcomes. Using a t-test, we examined if these 100 values were different from the 100 values of maximum cross-correlation coefficients produced for the same condition, taken as a measure of noise. In order to determine the ability of this procedure to detect changes in preferred orientation when shifting gaze across monitors, we simulated the expected changes by shifting randomly chosen tuning curves from the recorded neurons by a variable angle drawn from a discrete set of possible shifts (2, 4, 6, 8, and 10°). We then subjected the new data set (shifted) to the bootstrapping analysis described before. As shown in **Supplementary Figures S7A** and **B**, this simulation established that the bootstrapping analysis allowed us to detect significant shift angles down to 2°. We ran the bootstrapping analysis also on the original data, i.e. without the step of combining neighboring bins. As summarized in **Supplementary Figure S7C**, the result was similar, yet characterized by a lack of statistical power. Finally, we also confirmed that the level of noise characterizing the orientation tuning function of individual neurons did not influence our measurement of OUI (**Supplementary Fig. S7D**).

### Measuring the angular shift of the orientation preference (population analysis)

First, we normalized all neurons’ orientation tuning curves by scaling discharge rates for the various grating directions relative to the discharge rate for the preferred direction set to 1. Next, we rotated the orientation tuning functions obtained for the central monitor such as to align the preferred direction of a neuron with the head-centered horizontal (**Supplementary Fig. S8A; B** for M1 and M2). We then rotated a given neuron’s tuning curve for the eccentric monitors by the same amount. We kept all tuning curves at their original resolution, partitioning the range of possible orientations by bins of 2°. **Figure 2D** depicts the resulting normalized population orientation tuning functions for the three monitor positions on top of each other in a head-centered *FOR*. To measure the amount and the significance of the angular shift between two populations’ tuning functions for pairs of gaze directions, we used the 1D cross-correlation and bootstrapping procedures described before for single neurons.

### Measuring eye torsion

In order to determine the eye torsion associated with the three gaze directions, we analyzed high resolution images of both eyes, taken at 1 image/second as described in detail in Daddaoua and coworkers^20^. In brief, the torsional position was determined by identifying clearly visible landmarks on a small ink tattoo applied to the sclera (**Supplementary Table S2**). Next, a line was drawn between a distinct part of the landmark and the center of the pupil. By subtracting the orientation of the line for the central monitor from the orientation of the line for eccentric monitors, we could calculate the amount of torsion induced by oblique fixation. For more details see Daddaoua and coworkers^20^. Since we acquired hundreds of images in each session (for each eye one per second), the measurements were semi-automated through a pattern recognition approach: We started by determining the orientation of the line connecting the landmark and pupil in the first image as explained before. This orientation served as reference for subsequent images of the same eye for one experimental session and gaze direction. To measure the orientation in the subsequent images automatically, we first selected the pupil position on the reference image (*RI*) manually. Then, using 2D cross-correlation between the pupil in *RI* and the following image, we were able to detect the pupil position on all following images. This is possible since the resulting cross-correlation peaks when the pupil in the *RI* is aligned with the pupil in subsequent images. Using this same method, we measured the position of the landmark and calculated the orientation of the line connecting the landmark with the pupil center. We tested the accuracy and reliability of this semi-automatized method by comparing its performance with the one of an experienced human based on 450 pairs of images. The results were similar (**Supplementary Table S3**). In around 15% of the sessions we were able to acquire snapshots, yet without being able to use them for reliable torsional measurements because of too unfavorable lighting conditions. Hence, these sessions had to be discarded and the prevailing eye torsion estimated by resorting to the mean eye torsion for the given fixation condition, calculated from other sessions. This was justified in view of small variability of false torsion values for a given eccentricity as indicated by a 95% conf. b. of ± 0.04° (Fig. S3).

## Acknowledgments

We are grateful to Friedemann Bunjes and Peter W. Dicke for technical support. We would like to thank Nabil Daddaoua for his advice and comments. This work was funded by the German Research Foundation (Deutsche Forschungsgemeinschaft) grant number FOR 1847-A3 (TH 425/13–1), received by P.T. within the framework of the Research Unit “Primate Systems Neuroscience” and grant number EXC 307, Werner Reichardt Centre for Integrative Neuroscience (CIN), an excellence cluster grant at the University of Tübingen received by P.T. The authors declare no financial conflicts of interest.

## Author contributions

**P. T**. developed the conceptual framework of the research. **M. K**. and **P. T**. designed the study. **M. K**. collected the data and analyzed them. **M. K**. and **P. T**. discussed the experimental results and wrote the manuscript.

---

**Supplementary Figure S1.**
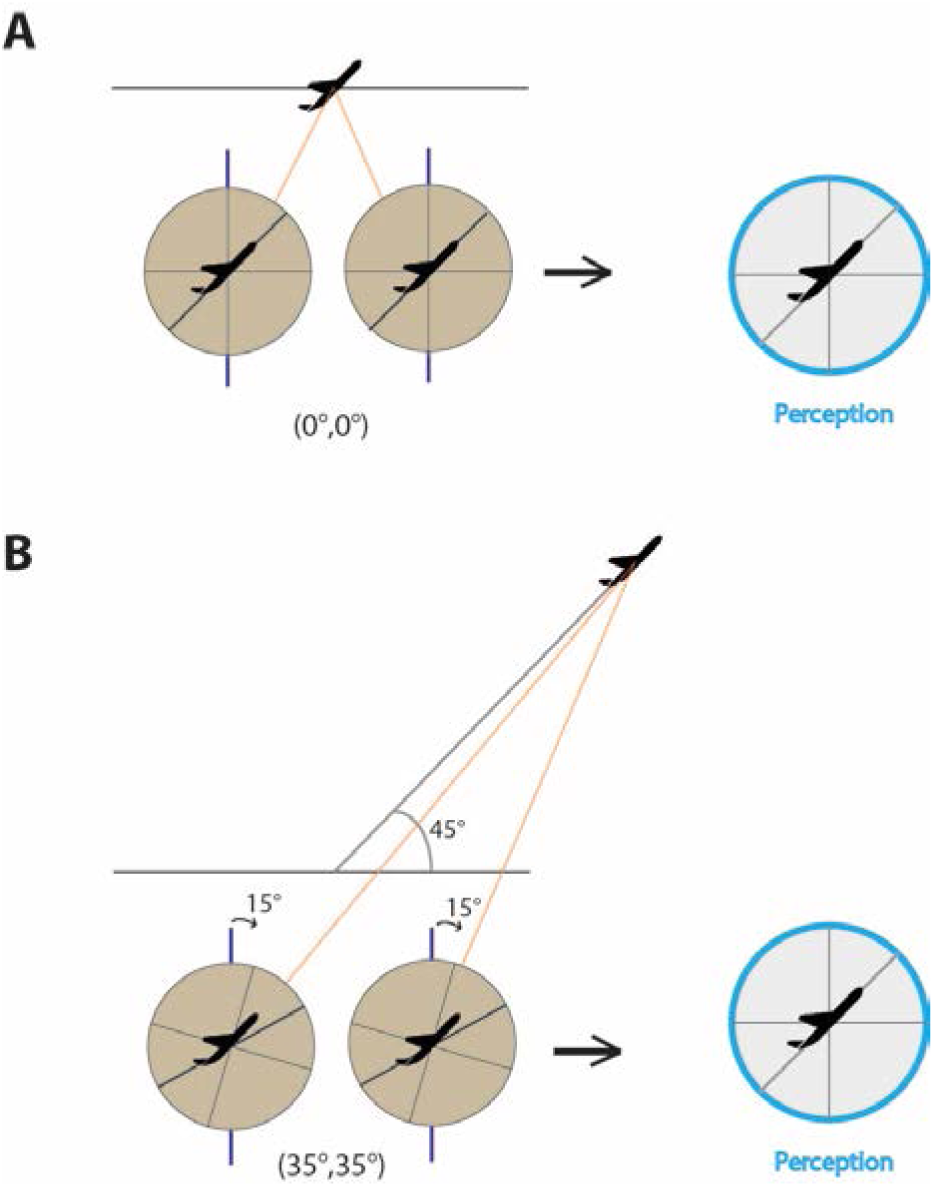
Illustration of the consequences of false torsion. **A** Both eyes are oriented straight ahead (0°, 0°) fixating an ascending airplane. The horizontal line marks the horizon**. B** A moment later the airplane has reached a more eccentric position relative to the head, forcing the fixating eyes to shift to the upper right (35°, 35°) with respect to straight ahead. Due to Listing’s law, this shift entails that the eyes rotate around the line of sight by approximately 15° (*“false torsion*”). We assume that the orientation of the airplane relative to the horizon is unchanged. Yet, as a consequence of *false torsion* its retinal image gets rotated in the opposite direction. However, this retinal rotation is not perceived. Rather the observer perceives the orientation of the airplane unchanged.

**Supplementary Figure S2.**
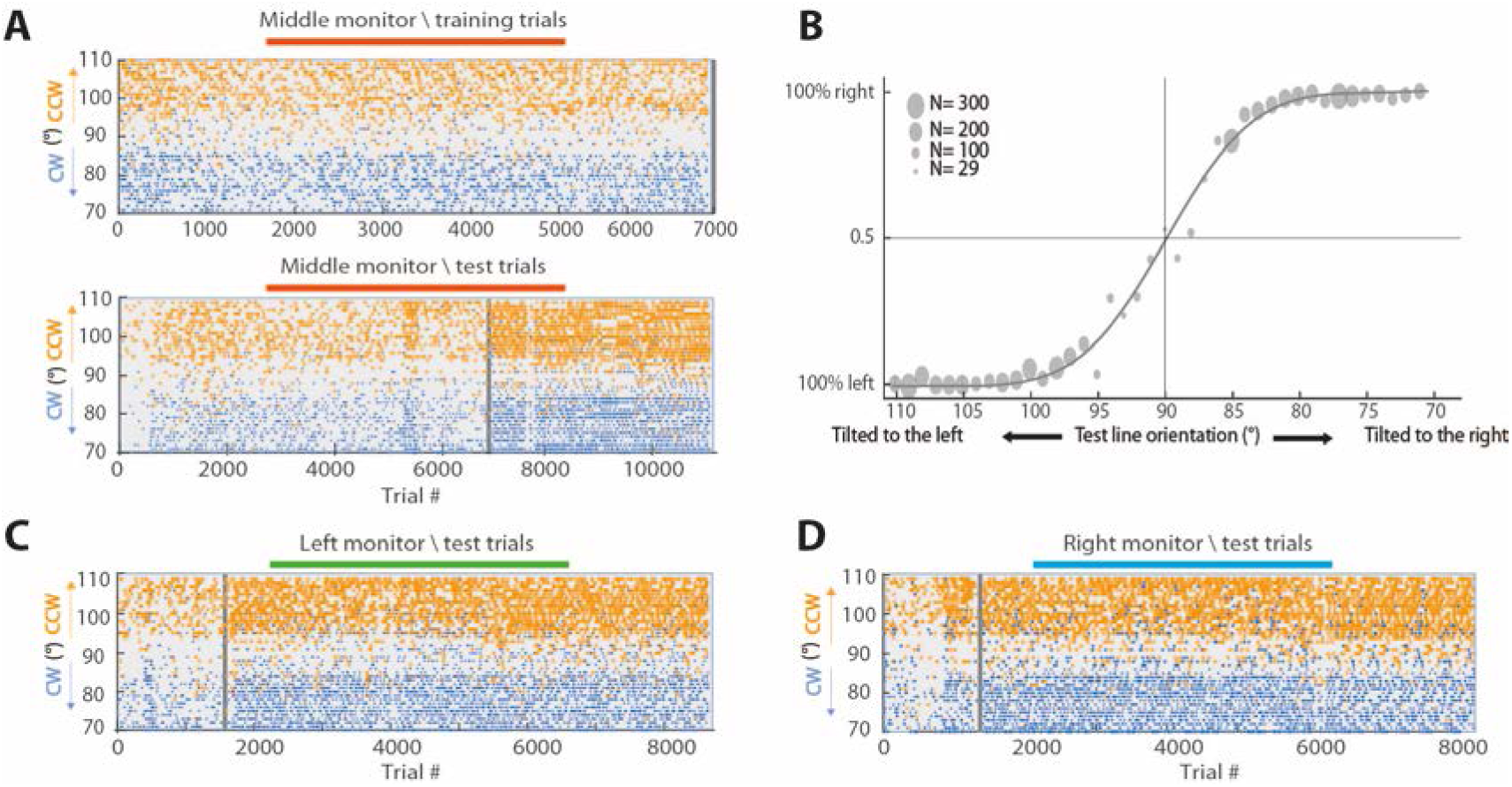
Summary of the reports of monkey M1 on perceived line orientation. **A**, **C** and **D** Raster plots of the monkey’s choices (“cw” vs. “ccw” line rotation relative to vertical) as a function of trial number. Each dot marks a perceptual decision and the choice is color-coded (blue=cw, orange=ccw). The y-axis specifies the true line orientation (90°=vertical, >90°=cw tilt, <90°=ccw tilt) on the monitor and the x-axis the trial number. **A, upper panel** All training trials for the central monitor, for which M1 had to indicate the line orientation in the presence of a reference line indicating the vertical. These trials were performed in the first weeks. From trial number 7,000 onwards, marked by the gray vertical line, the presentation of a reference line could be stopped as the monkey had understood the task. **A, lower panel** All test trials for the central monitor in which a reference line was not present (in 10–20% of the cases before trial 7,000 and consistently thereafter)**. B** Psychometric curve based on data from (**A**). The diameter of the dots quantifies the number of *cw* or *ccw* decisions for a given line orientation. The PES derived from the curve corresponds to 90.33° (conf.b 90.29, 90.37). **C** and **D** Decisions on test trials for the upper left monitor and upper right monitor respectively. As in (**A**) the gray vertical line separates early test trials, which were still presented interleaved with training trials in which the reference line was available and later trials without reference line. Note fact that the density of test trials in (**A**, **C** and **D**) is relatively low in the beginning early in the experiment (relative to the grey vertical line) as in this phase they were mixed with training trials. Note as well that the frequency of test line tilts close to the true vector orientation was kept low (10–15%) in order to ensure that the monkey’s concept of vertical would not be compromised by too many close-to-threshold decisions.

**Supplementary Figure S3.**
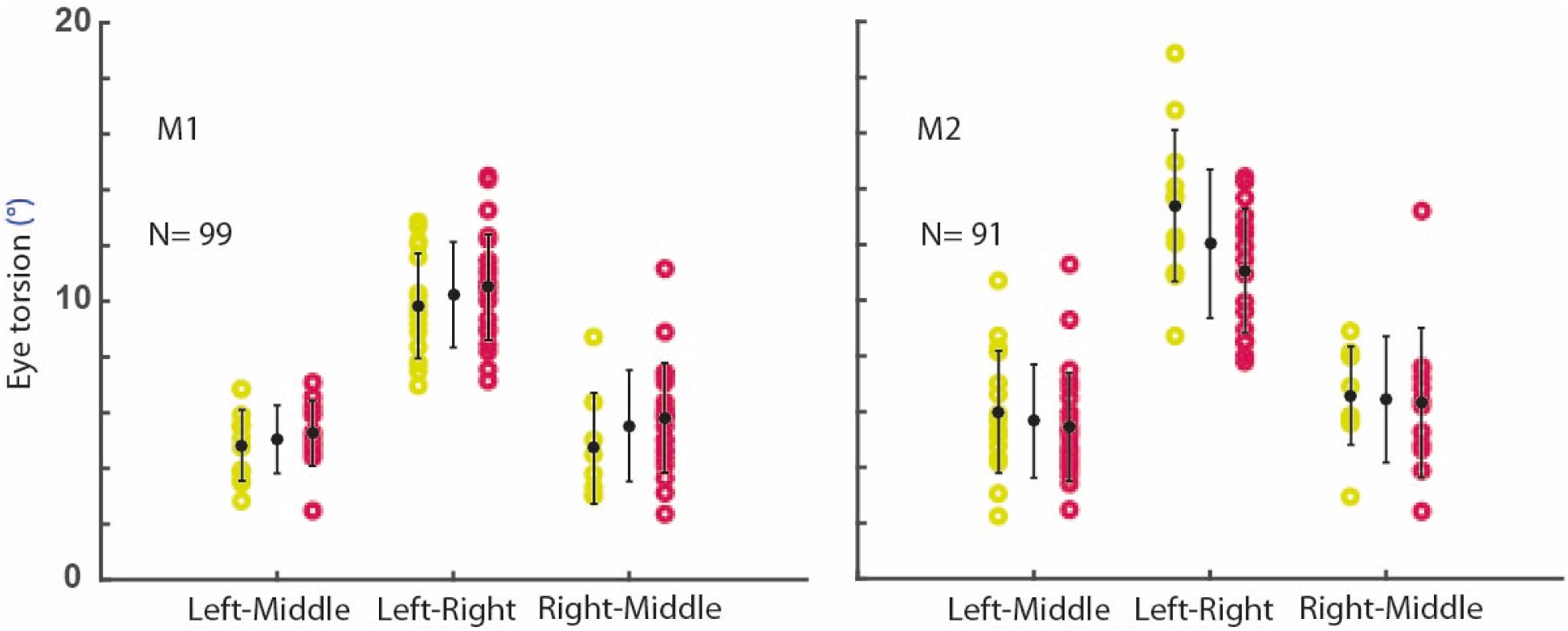
Eye torsion dependency of gaze direction. Summarizes the changes in torsion of the two eyes (right eye: yellow, left eye: red) for the different gaze directions compared. Data for monkey M1 is shown on the left, data for monkey M2 on the right. Each data point represents the difference between static levels of eye torsion maintained while assessing the features of an individual neuron for the two gaze directions compared. The error bars indicate standard deviations.

**Supplementary Figure S4.**
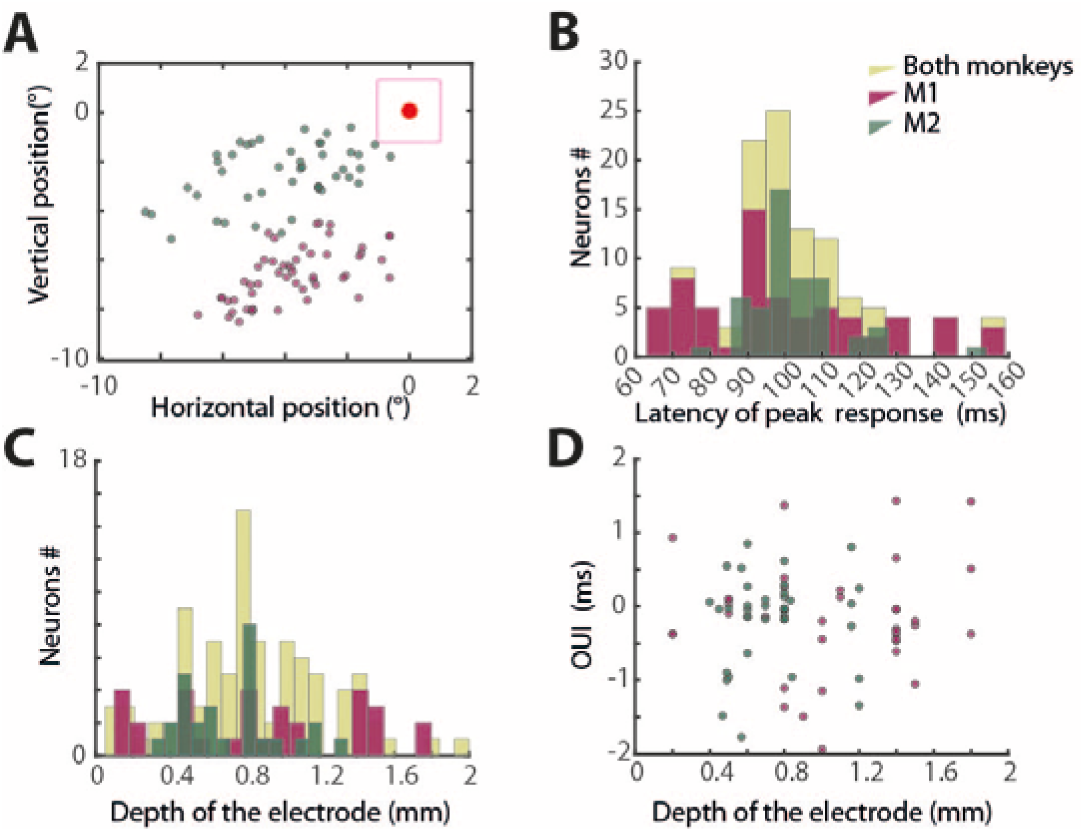
RFs locations, response latencies and recording depths of the studied neurons. **A** Location of RFs of neurons from M1 (purple) and M2 (dark green). The red dot marks the position of the fovea**. B** Distribution of peak response latencies of the V1 neurons tested as derived from the reverse correlation analysis of RF measurements (see Online Methods). The distributions for the two monkeys are shown in red (M1) and green (M2) and the pooled distribution in yellow. The latencies of visual responses were on average 99.61 ms ±25.3 standard deviation (stdv) from M1 and 102.3 ms ±12.2 stdv for M2. **C** Distribution of recording depths of individual neurons. The red and green distributions indicate neurons from monkeys M1 and M2, which could be tested for at least two monitors. The yellow distribution is the composite of the two, complemented by neurons, which could be tested only for one monitor. **D** Plot of OUI of individual neurons as function of recording depth. Note that there is no significant correlation between the two variables (r=-0.062, p=0.57).

**Supplementary Figure S5.**
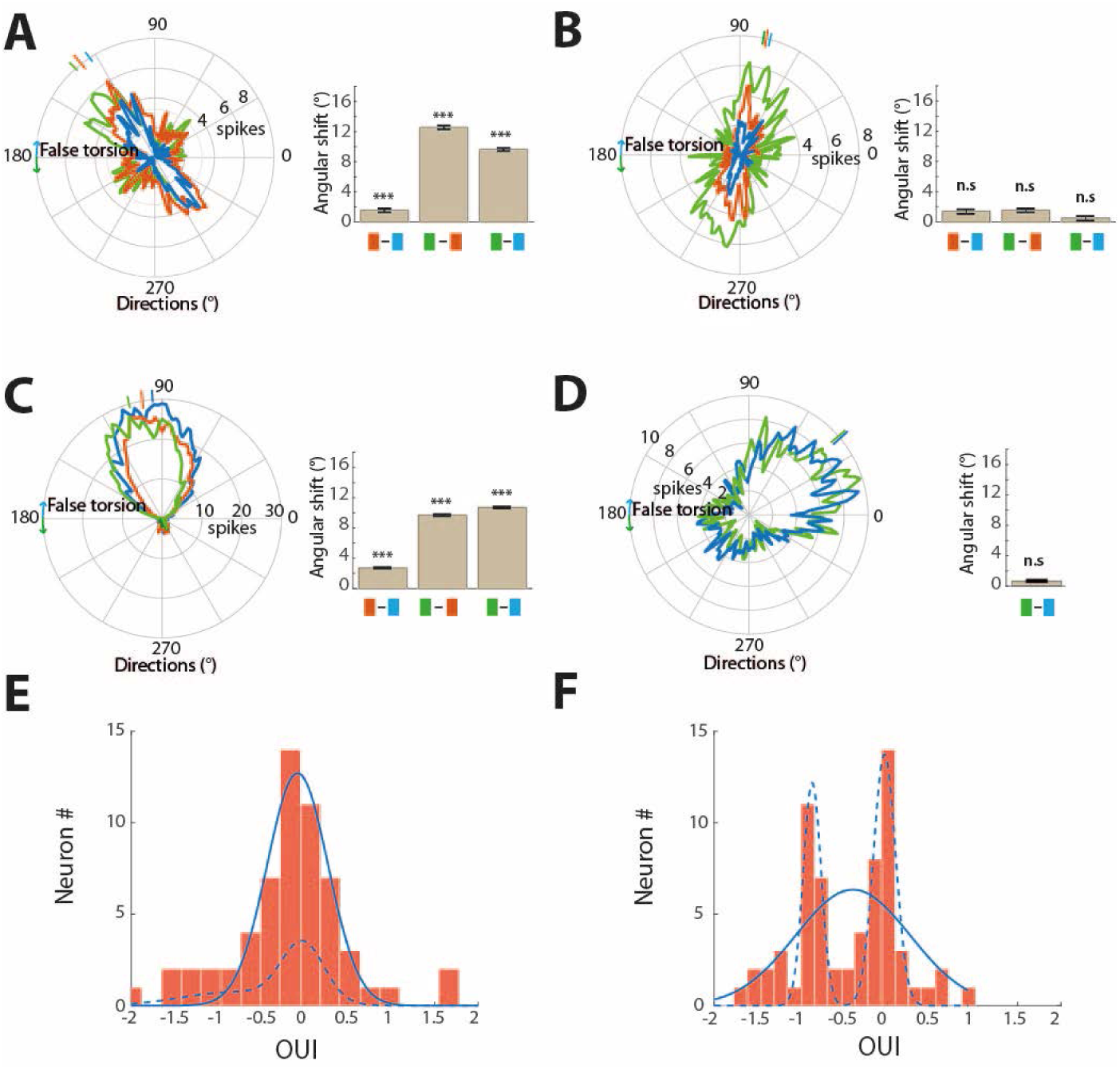
Exemplary V1 neurons encoding visual orientation in different FOR. **A, B, C** and **D** Orientation tuning curves of four individual neurons plotted in a head-centered FOR. The bar charts next to each tuning curve quantify how much the preferred orientation changed in head-centered coordinates when shifting fixation from one monitor to another one. Color code: green, data for upper-left monitor, blue for upper-right one and brown for the central one. The orientation preferences of the neurons in (**A**) and (**C**) exhibit substantial shifts, in some conditions approaching the changes in eye torsion, indicating that these neurons preferred a retina-centered FOR. Conversely, the tuning curves of the neurons depicted in (**B**) and (**D**) were more stable in head-centered coordinates, suggesting a preference for head-centered coding. E and F plot the distribution of OUIs for all neurons from M1 and M2 respectively.

**Supplementary Figure S6.**
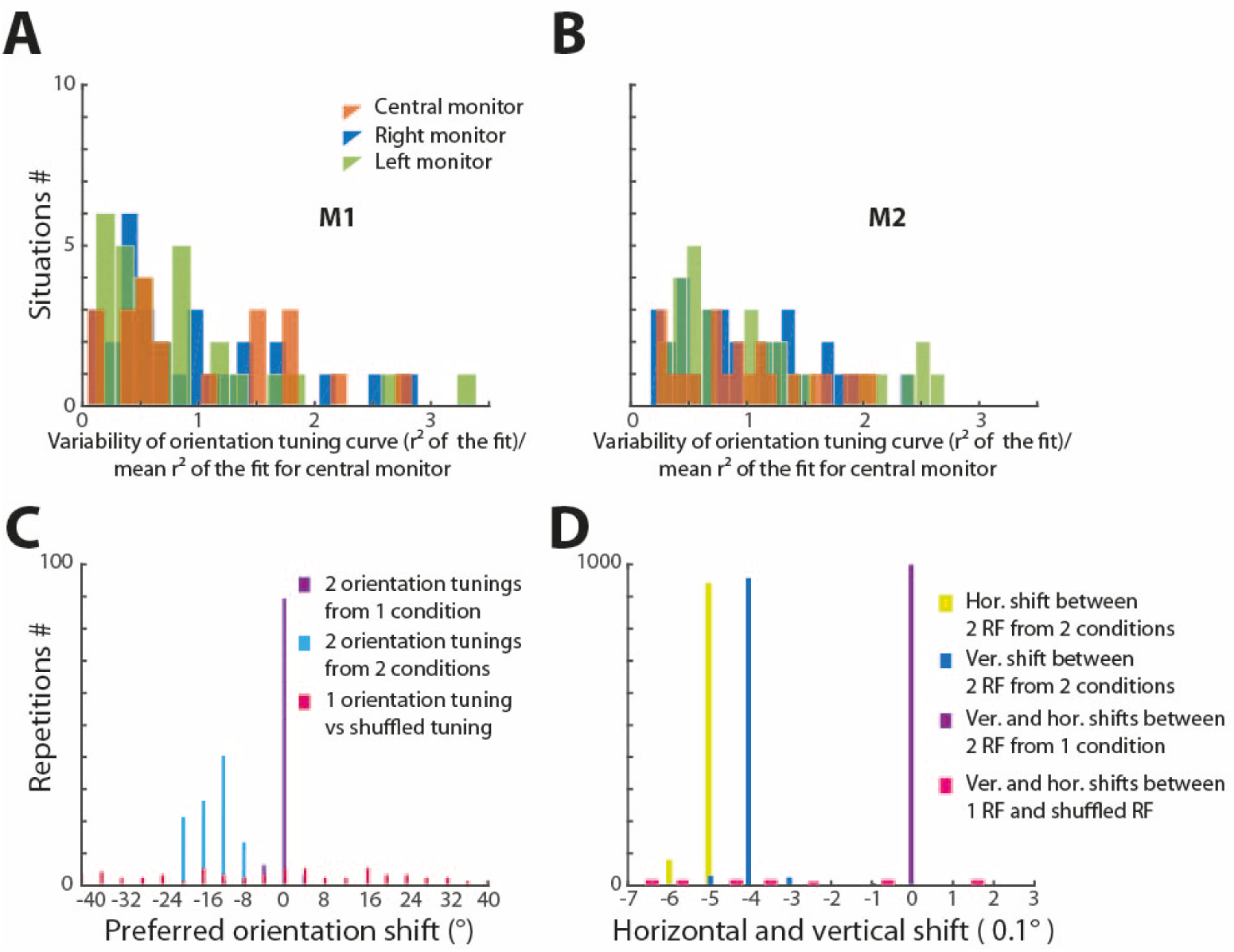
Constant variability in the orientation tuning across the three monitors. **A** and **B** Histograms of the normalized r² representing the goodness of the orientation tuning curve fit (see Online Methods) for the three monitors for monkey M1 **(A)** and M2 **(B)**. The histograms do not reveal a significant influence of gaze direction on the goodness of fit, i.e. the variability of the orientation tuning curves (1-way ANOVA with the factor gaze direction, p=0.89 for M1 and p=0.85 for M2. **C** and **D** illustrate the reliability of our method of measuring the shifts of orientation preference and RFs by cross-correlating bootstrapped populations from neural responses. **C** plots the repetitive measurement of the preferred orientation shift based on bootstrapping and subsequent cross-correlations (see Online Methods). The shift between two orientation tuning curves of one neuron measured for the upper right and the upper left monitors (blue) is significant. This shift is significantly different from the shift predicted when comparing two sets of bootstrap populations generated from data for a neuron tested with stimuli on one monitor only (purple). The same method fails to yield any reliable shift between one orientation-tuning curve and itself after shuffling (noise) (light purple). **D** plots the transitional shift of RF in a similar manner to (**C**). Note that the horizontal and vertical shifts are reliably measured.

**Supplementary Figure S7.**
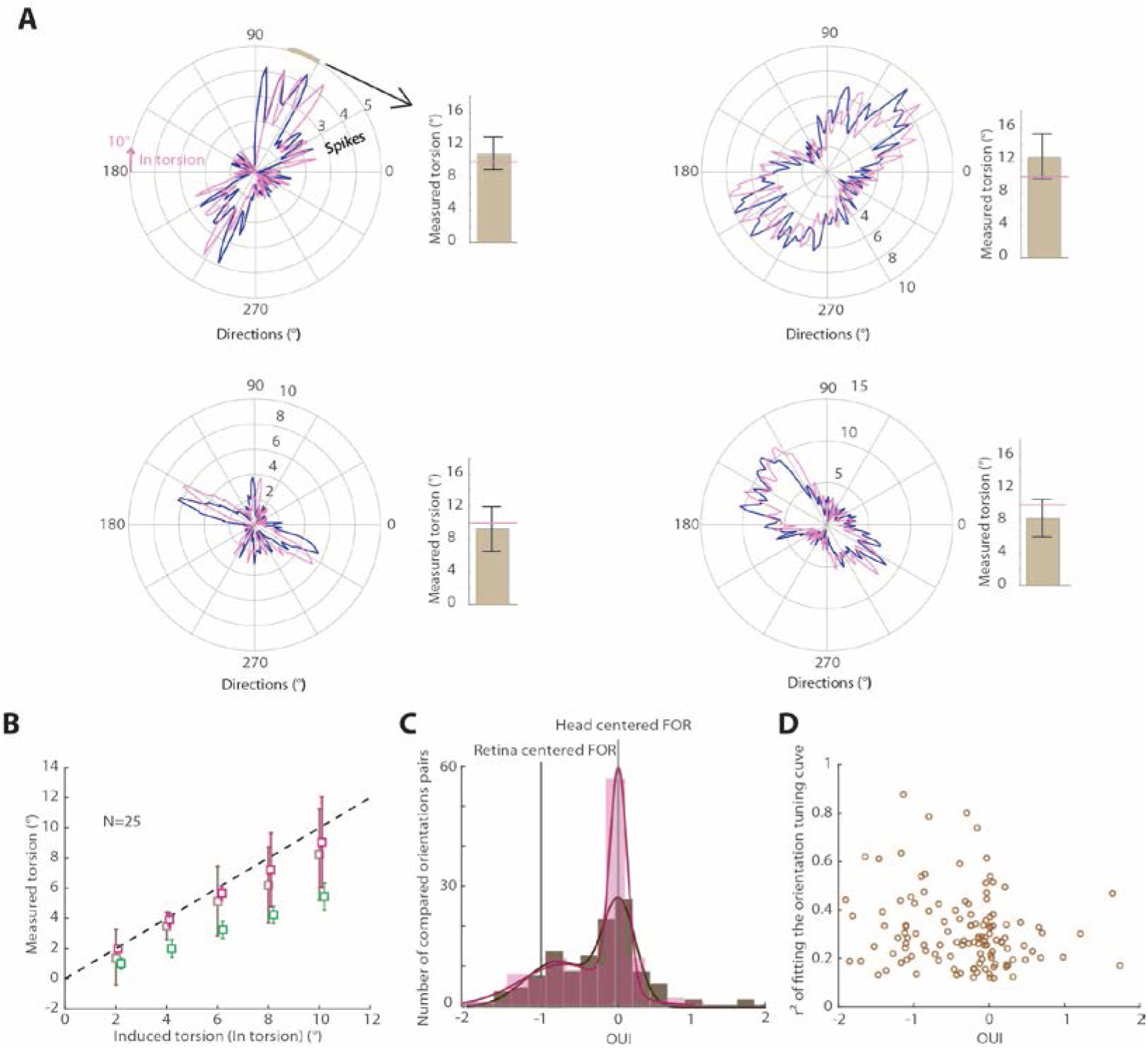
Examination of the sensitivity of our measurement of orientation preference changes. **A** The dark blue lines are the orientation tuning curves of four exemplary V1 neurons plotted in head-centered polar coordinates. The purple curves are the tuning curves after 10° clockwise rotation. The bar plots give the amount of orientation preference changes, determined by the bootstrapping-cross-correlation analysis described in Online Methods. The error bars represent RMSE and the horizontal purple line indicates a 10° rotation. Note that the measured rotation is very close to the actual imposed rotation. **B** Comparison of the results of different ways to measure angular changes of preferred orientation imposed by rotating original tuning curves by known amounts. Data obtained with the bootstrapping-cross-correlation approach underlying the analysis presented in the main text is shown in brown, results obtained from cross-correlating the original tuning curves in purple and results derived from a cross-correlation of the orientation functions obtained by fitting the original tuning curves19 in green. This comparison is based on 25 neurons with tuning curves representing the full spectrum of different degrees of tuning sharpness. Note that in principle all three approaches worked, yet the one relying on the original tuning curves substantially underestimated the true amount of orientation preference change. **C** Comparison of the two histograms of OUIs for all recorded neurons. OUIs values obtained based on cross-correlating mean tuning curves without bootstrapping are shown in lighter color and those based on repeated cross-correlations of bootstrapped populations of the measured tuning curves are depicted in darker color (see Online Method). Note that the two histograms are very similar (p=0.64, Wilcoxon rank sum). **D** Plot of the goodness of the fit of the (r²) orientation tuning curves to the orientation-tuning model proposed by Wörgötter and Eysel19, against OUI. Note that the quality of the fit did not affect our measure of OUI (r=-0.06, p=0.47).

**Supplementary Figure S8.**
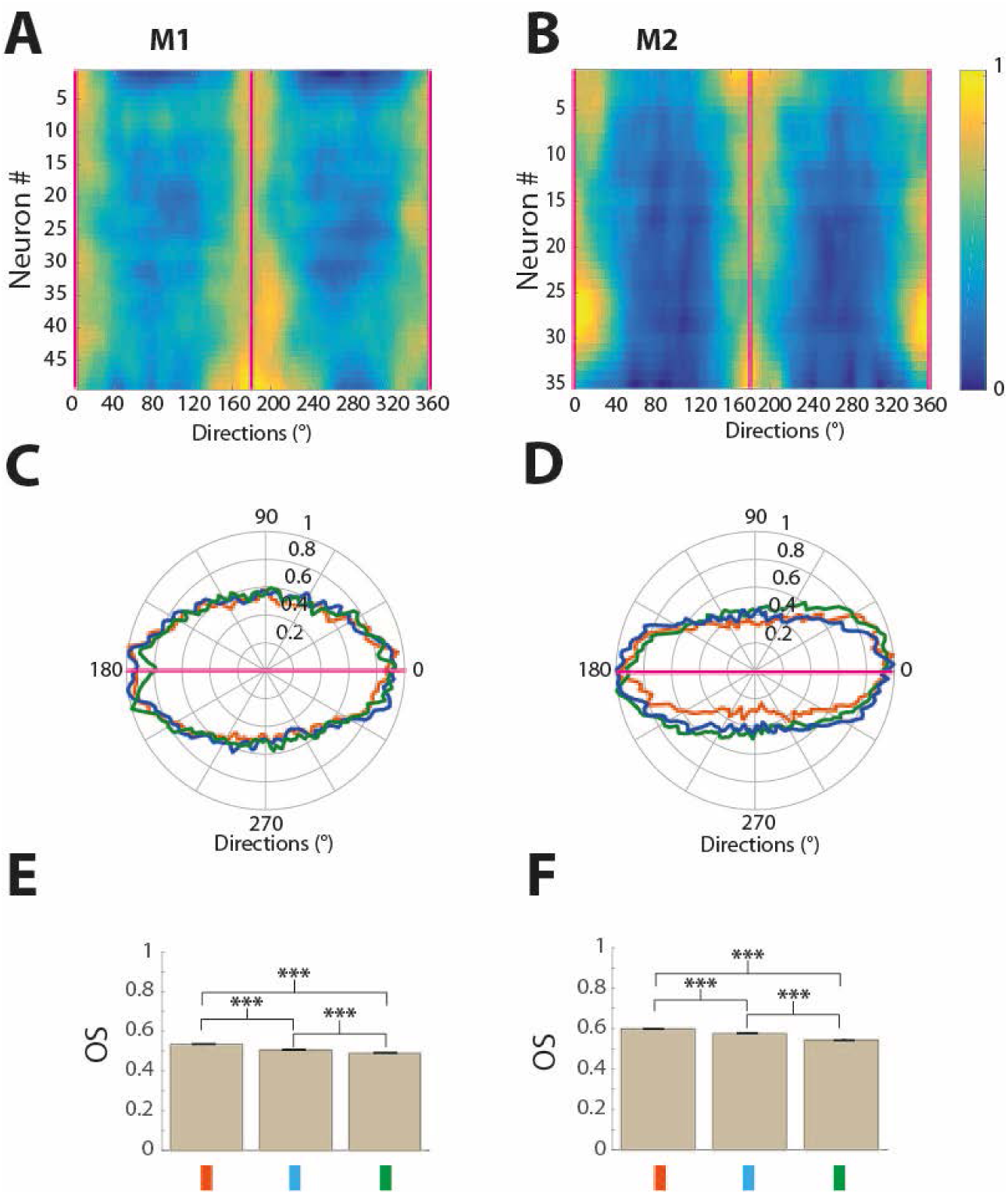
Population orientation tuning curves for the two monkeys (left column M1, right column M2). **A** and **B** Color maps representing the orientation preferences of all neurons for the central monitor after alignment of their respective preferred orientation with the horizontal. Firing rates are normalized relative to the firing rate for the preferred orientation set to 1 (indicated by yellow; deep blue =0). **(A)** neurons from M1 and **(B)** from M2. **C** and **D** depict the population tuning curves for the three monitors for M1 and M2 with data for the eccentric monitor aligned with respect to the central one (see Online Methods for details). Data for upper-right monitor in blue, for the upper-left one in green and red for the central monitor. **E** and **F** Population means and se of orientation selectivity indices OS for the three gaze directions (color code as before). Data for M1 on the left, for M2 on the right. The asterisks mark significant pairwise comparisons of the means (t-test, ***= p<0.001).

**Supplementary Figure S9.**
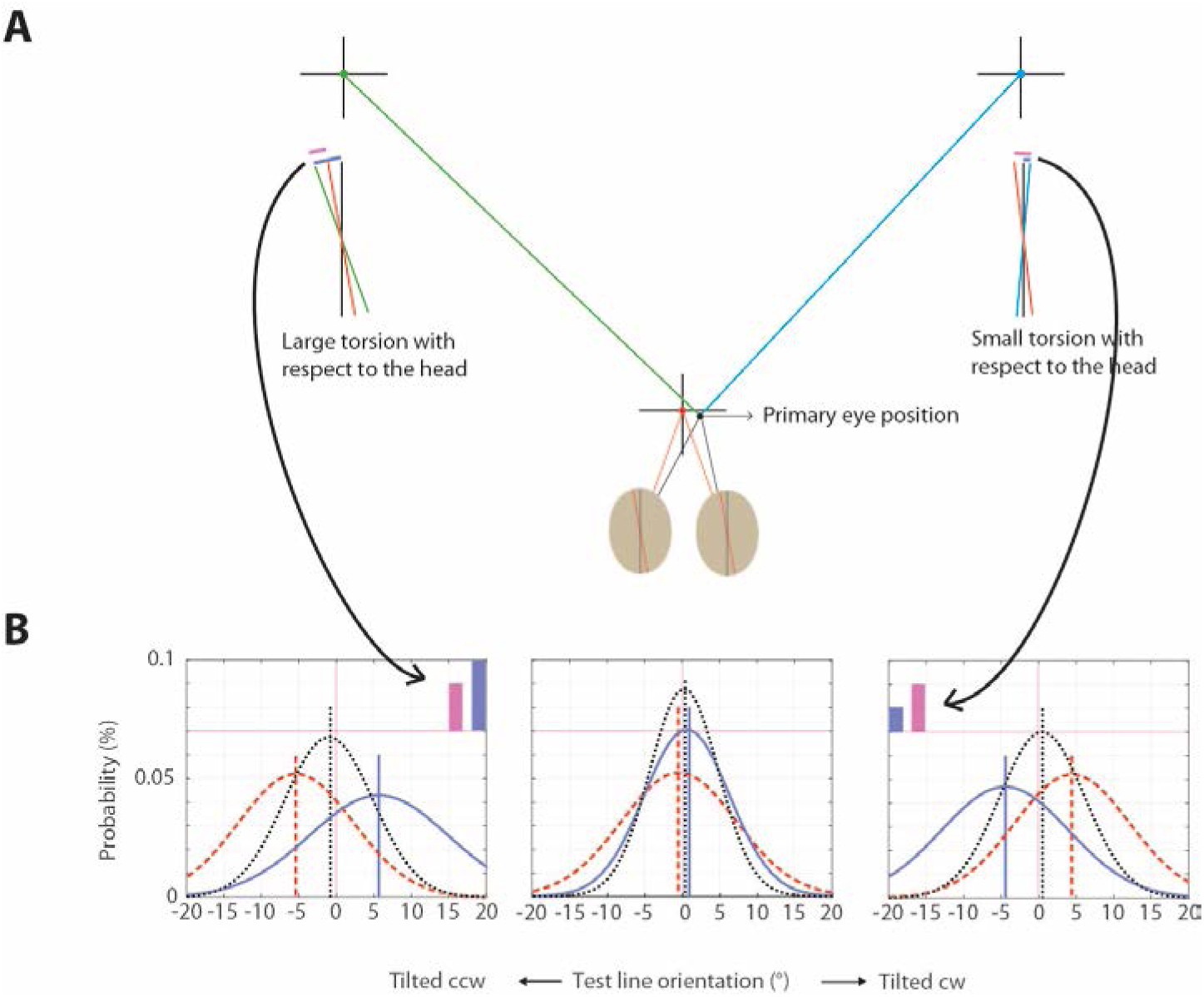
Bayesian integration model of orientation perception. As explained in the Online Methods part, the model assumes that orientation judgments are based on the integration of information on retinal orientation provided by visual orientation filters and an eye torsion prior that encodes torsion in a linear format. **A** presents an illustration of three distinct scenarios, namely the eyes looking straight ahead and exhibiting a subtle torsion bias in a 0.53° *ccw* direction, alternatively the eyes deviating a little bit in a direction inducing a small amount of *false torsion* that aligns the retinal meridian with the world vertical line and finally, the eyes adopting a fairly eccentric orientation causing a much larger amount of *false torsion*. Because of the assumed eye torsion bias the retinal vertical meridian is rotated *ccw* relative to world vertical for gaze straight ahead. And because of the bias, the eye torsion prior will be larger for gaze shifts to the upper left (green) than to the upper right monitor (blue). **B** The three figures plot functions that give the probability of perceiving a certain orientation (x-axis) as vertical (SVV) based on which signals contribute to perception —. i.e., the probability function arising from just the retinal signal (dashed red line) predicts the world vertical line to be aligned always with the retinal meridian, whereas the ones based on the eye torsion prior only predicts it shifted by the same amount as the latter but with opposite direction (blue line) and finally, the resulting estimate of the SVV based on integrating both signals (dotted black line) locates it in between the three gaze directions distinguished in **(A)**. The SVV estimate is given by maximum likelihood fit of the psychophysical data (see Online Methods). Note that the position of the peak of the estimated SVV is not predicted by gaze direction whereas its width increases with eye torsion. In the case of the torsion prior signal, the gaze direction changes the peak position of the predicted world vertical orientation and the width of the probability function, based on a standard deviation of **σ**=a0+a1∙***torsion***. Fitting the model with the psychophysical data results with the parameters **σ_0_**=5.3, **σ_1_** =0.72, eye torsion bias during straight head fixation= 0.53° *ccw* and retinal signal noise having standard deviation of **σ_r_**= 7.7. Note the purple bars on the left and right panels. The light purple bar represents the difference in *false torsion* between the eccentric gaze directions and straight-ahead direction, which was similar for both left and right monitors. However, due to the torsional bias, the model predicts a larger amount of false torsion for the left monitor with respect to the world vertical as compared with the right monitor.

**Supplementary Figure S10.**
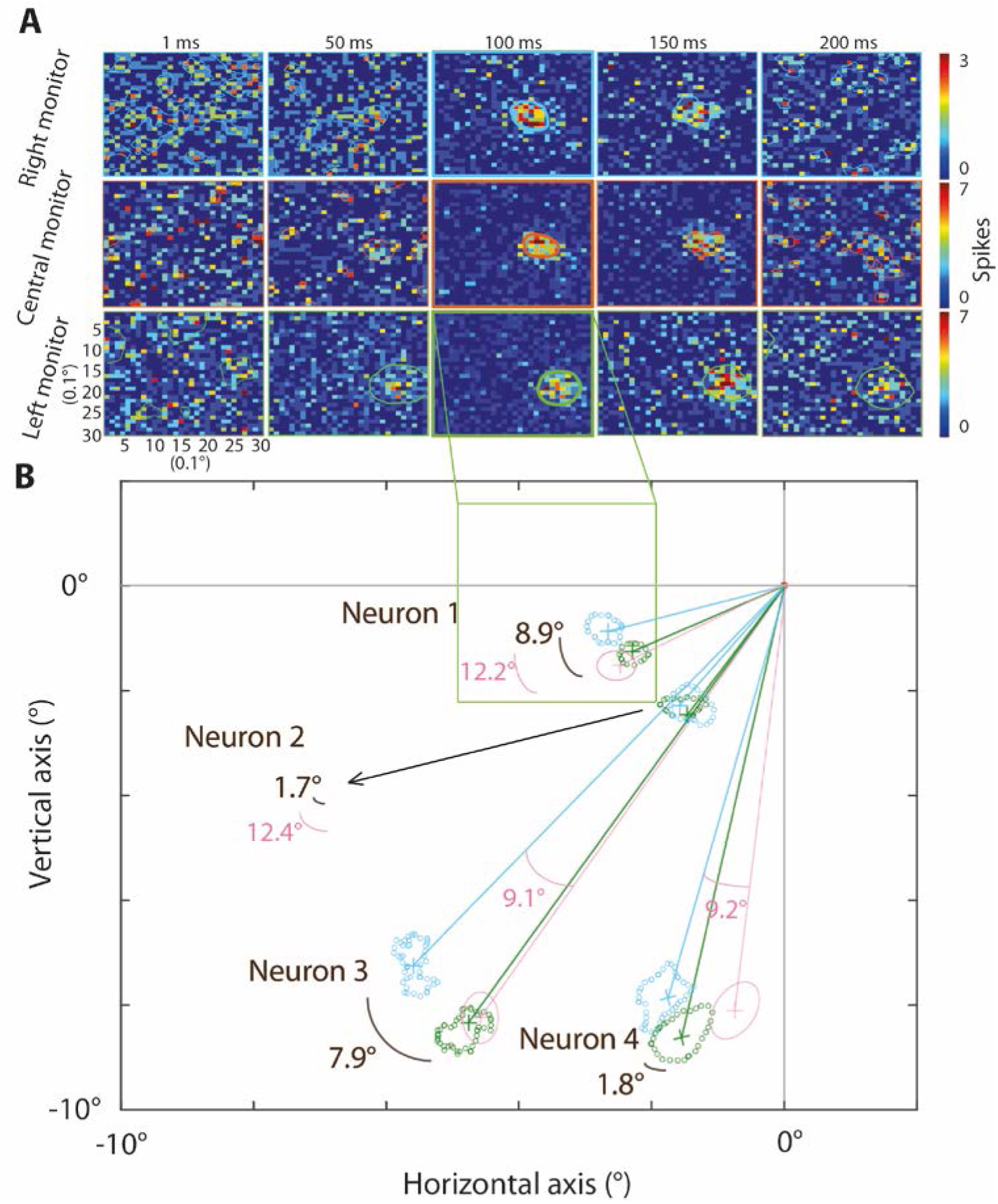
Plot of false torsion impact on the RF location in head-centered FOR. **A** RF maps of an exemplary V1 neuron at different times relative to stimulus onset (t=0 ms) as revealed by reverse correlation. Plots are in head-centered coordinates and the three rows depict the maps obtained for the three gaze directions (i.e. fixation on the upper left, the central and the upper right monitor). The maps highlighted by bold colored contour lines represent the peak responses. The peak response and the time of its occurrence were determined by summing all spikes along the y-dimension, giving a 1-dimensional vector of elements presenting the number of spikes along the x-dimension. The time relative to stimulus onset yielding the x-vector with the largest peak was then taken as the time of the peak response. The bold colored contour lines demarcate the boundaries of the RFs at the time of the peak response. The contours obtained for fixation on the two eccentric monitors are reproduced in the visual field map with respect to the fixation point depicted in **(B)**. **B** This map compares the RFs of neuron 1 and three others in a head-centered FOR for fixation on the two eccentric monitors. RFs for gaze on the upper right monitor are shown in blue, those for gaze on the upper left in green. In addition, the RFs positions for gaze on the upper left monitor to be expected if neurons used a purely retina-centered FOR are indicated by schematic elliptic RF boundaries (pink). The dark brown arc segments display the effective angular shift of RFs in head-centered coordinates (numbers attached), whereas the pink arc segments indicate the difference in *false torsion*.

**Supplementary Figure S11.**
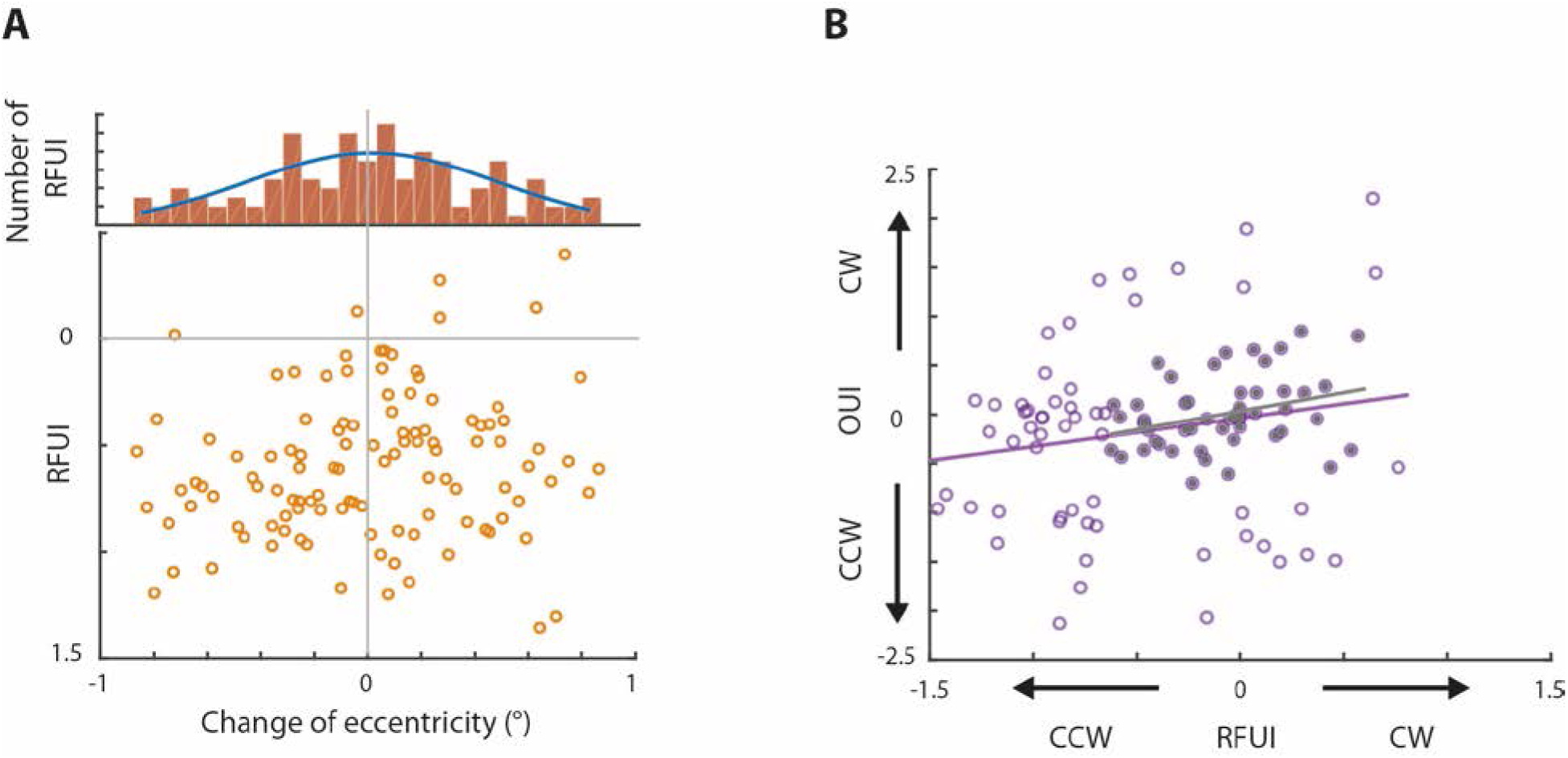
A plot of the relationship between the RF updating index RFUI and changes of RF eccentricity. **A** The histogram plot shows that changes of RF eccentricity cluster around zero (p=0.89, t-test) without relationship to the changes of the RFUI (r=0.15, p=0.1). In other words, only the angular position of RFs but not the eccentricity of RF change with *false torsion*: This is consistent with the expectation that the observed angular RF position changes are due to *false torsion*. **B** Plot of the amount and direction of orientation preference shifts against the amount and direction of RF shift of the same neuron. The plot is based on subsets of neurons that have RFUIs and OUIs either within ±2.5 standard deviation (stdv) of the population RFUIs and OUIs (open symbols) or ± 1.0 stdv (full symbols). The correlation between the two variables is in any case significant but strengthened by constraining the data set to neurons whose indices are closer to the respective means (±2.5 stdv subset: r=0.19, p=0.044, n=109; ±1.0 stdv subset: r=0.3, p=0.027 n=55). The significant correlations are consistent with the view that both indices reflect adjustments to *false torsion*.

## Supplementary tables

**Supplementary Table 1.** Means and standard deviations of fixation positions on all three monitors for M1 and M2. The values are based on periods of fixation during periods of testing all neurons considered for later analysis of torsion-dependent changes.

|  | Monkey M1 |  |  |  |  |  | Monkey M2 |  |  |  |  |  |
| --- | --- | --- | --- | --- | --- | --- | --- | --- | --- | --- | --- | --- |
| Monitor | left |  | central |  | right |  | left |  | central |  | right |  |
| Eye position / measurement | mean | stdv | mean | stdv | mean | stdv | mean | stdv | mean | stdv | mean | stdv |
| x | -0.059 | 0.15 | -0.074 | 0.19 | -0.12 | 0.17 | 0.19 | 0.31 | -0.1 | 0.23 | -0.016 | 0.1 |
| y | -0.061 | 0.17 | -0.07 | 0.21 | -0.049 | 0.22 | -0.15 | 0.27 | -0.18 | 0.24 | -0.025 | 0.024 |

**Supplementary Table 2.** Reliability of using landmarks in eye snapshots in order to assess eye torsion changes. Means and standard deviations of the orientation of the line connecting the chosen landmark with the pupil center based on n=30 repetitions, using the same eye snapshots, repeated on 15 randomly chosen photographs. The mean line orientation is taken as 0° orientation. The smallness of the standard deviations documents the high reliability of the method.

|  | Monkey M1 |  | Monkey M2 |  |
| --- | --- | --- | --- | --- |
|  | mean (°) | stdv (°) | mean (°) | stdv (°) |
| Left eye | -2.3e-15 | 0.32 | -8.6e-17 | 0.24 |
| Right eye | 4.7e-16 | 0.31 | 6.4e-16 | 0.27 |

**Supplementary Table 3.**
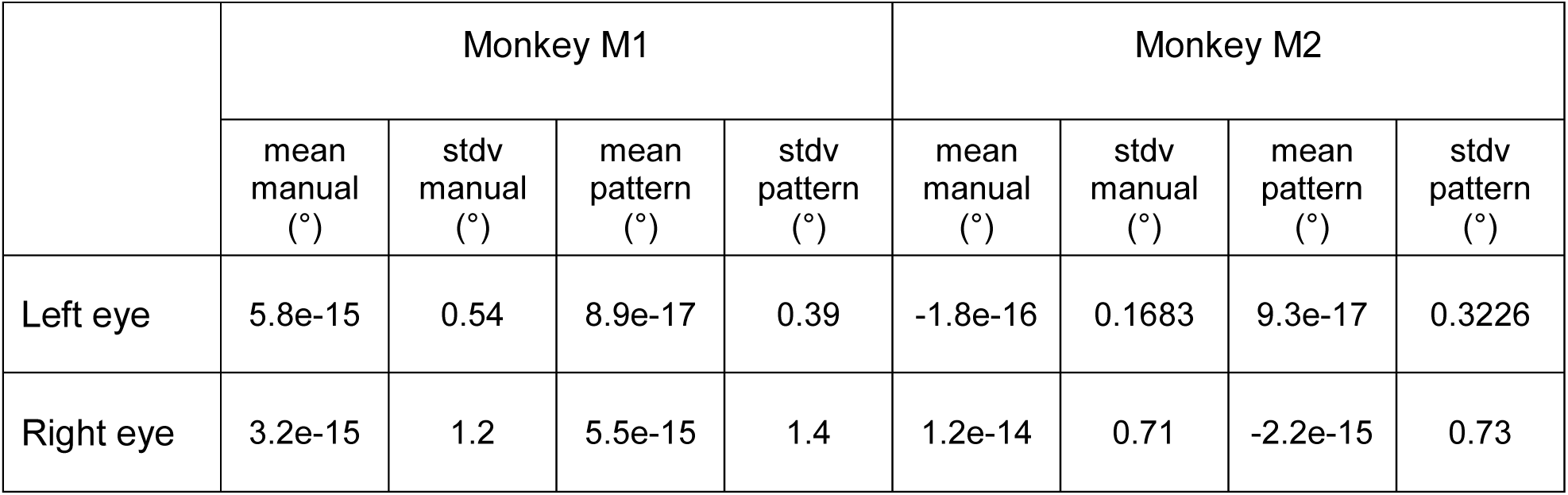
Comparison of manual analysis of landmark orientation in eye snapshots with a semiautomatic approach involving pattern recognition (see Online Methods for details). The means and deviations of eye torsions are based on 450 snapshots from one experimental session with mean eye torsion defined as 0° orientation. The comparison documents using the pattern recognition-

